# The landscape of *Arabidopsis* tRNA aminoacylation

**DOI:** 10.1101/2024.09.09.612099

**Authors:** Luis F. Ceriotti, Jessica M. Warren, M. Virginia Sanchez-Puerta, Daniel B. Sloan

## Abstract

The function of tRNAs depends on enzymes that cleave primary transcript ends, add a 3′ CCA tail, introduce post-transcriptional base modifications, and charge (aminoacylate) mature tRNAs with the correct amino acid. Maintaining an available pool of the resulting aminoacylated tRNAs is essential for protein synthesis. High-throughput sequencing techniques have recently been developed to provide a comprehensive view of aminoacylation state in a tRNA-specific fashion. However, these methods have never been applied to plants. Here, we treated *Arabidopsis thaliana* RNA samples with periodate and then performed tRNA-seq to distinguish between aminoacylated and uncharged tRNAs. This approach successfully captured every tRNA isodecoder family and detected expression of additional tRNA-like transcripts. We found that estimated aminoacylation rates and CCA tail integrity were significantly higher on average for organellar (mitochondrial and plastid) tRNAs than for nuclear/cytosolic tRNAs. Reanalysis of previously published human cell line data showed a similar pattern. Base modifications result in nucleotide misincorporations and truncations during reverse transcription, which we quantified and used to test for relationships with aminoacylation levels. We also determined that the *Arabidopsis* tRNA-like sequences (t-elements) that are cleaved from the ends of some mitochondrial mRNAs have post-transcriptionally modified bases and CCA-tail addition. However, these t-elements are not aminoacylated, indicating that they are only recognized by a subset of tRNA-interacting enzymes and do not play a role in translation. Overall, this work provides a characterization of the baseline landscape of plant tRNA aminoacylation rates and demonstrates an approach for investigating environmental and genetic perturbations to plant translation machinery.

## INTRODUCTION

Transfer RNAs (tRNAs) play a pivotal role in the central dogma of molecular biology by recognizing codons in messenger RNAs (mRNAs) and delivering the corresponding amino acid during protein synthesis. The expression and maturation of tRNAs depends on interactions with numerous specialized enzymes (Phizicky and Hopper 2010). Precursor tRNA transcripts are processed by the endonucleases RNase P and RNase Z to remove their 5′ leader sequence and 3′ extension, respectively (Hartmann et al. 2009). The mature 5′ and 3′ ends then base pair to form the acceptor stem with a 3′ overhanging nucleotide known as the discriminator base, which is followed by a 3′ trinucleotide CCA sequence. This CCA tail can be genomically encoded but is more often incorporated post-transcriptionally by a CCA-adding tRNA nucleotidyltransferase (CCAse), which plays an important role in tRNA quality control (Betat and Mörl 2015). The maturation of tRNAs also involves extensive post-transcriptional base modifications, which can be essential for proper tRNA folding, stability, codon-anticodon recognition, and charging by aminoacyl-tRNA synthetases (aaRSs) (Motorin and Helm 2010; Suzuki 2021; Giegé and Eriani 2023). Aminoacylated tRNAs deliver their amino acid to the ribosome for addition to the growing polypeptide chain during protein synthesis, after which they can cyclically repeat the process through multiple rounds of aminoacylation.

Availability of aminoacylated tRNAs is, thus, essential for sustaining protein synthesis. The levels of aminoacylation in the cellular tRNA pool are also environmentally responsive and play important signaling roles (Dittmar et al. 2005; Subramaniam et al. 2013; Raina and Ibba 2014; Avcilar-Kucukgoze et al. 2016). Being able to measure the relative abundance of aminoacylated and uncharged tRNAs is vital for understanding the associated regulatory mechanisms.

In addition to canonical tRNAs, there are many transcripts with tRNA-like sequences that can be recognized by the same suite of enzymatic machinery. For example, the maturation of human mascRNA (MALAT1-associated small cytoplasmic RNA) depends on a tRNA-like structure that recruits RNase P and RNase Z for cleavage of a long noncoding RNA (lncRNA) precursor transcript. However, mascRNA is sufficiently divergent in the structure of its D stem and anticodon region that it is not aminoacylated by aaRSs and does not interfere with translation (Skeparnias et al. 2024). Another example is the tRNA-like structure involved in the replication, translation, and genome encapsidation of the brome mosaic virus in plants. In this case, however, recognition and aminoacylation by a host aaRS is necessary for the full functionality of the viral element (Bonilla et al. 2021).

Plant mitochondrial genomes contain another class of tRNA-like sequences (t-elements) that are homologous to tRNA genes and encode transcripts predicted to retain typical tRNA-like secondary structures (Hanic-Joyce et al. 1990; Forner et al. 2007). Because t-elements often occur at the 5′ or 3′ ends of plant mitochondrial genes, recognition and cleavage of these structures by RNase P and RNase Z can define the ends of mature ribosomal RNA (rRNA) and mRNA transcripts (Forner et al. 2007; Warren et al. 2021b). Initial characterization of t-elements also indicated that they could act as CCAse substrates *in vitro*, but they were not detectable *in vivo*, suggesting they were unstable and rapidly removed (Hanic-Joyce et al. 1990). However, subsequent investigation found that t-element transcripts are at least transiently present *in vivo* and can carry post-transcriptionally added 3′ nucleotides that are not genomically encoded (Forner et al. 2007). Although this combination of findings suggests the action of a CCAse, a typical CCA motif was rarely observed in the analysis of *in vivo* transcripts. Indeed, in the most extensively studied example (the t-element that is transcribed at the 5′ end of *cox1*), sequenced transcripts had various strings of Cs and/or As but none of them had a standard CCA tail (Forner et al. 2007). These findings raise multiple alternative hypotheses: (1) the observed post- transcriptional addition of 3′ nucleotides to t-elements *in vivo* is not mediated by a conventional CCAse, (2) CCAse activity is less specific when not acting on canonical tRNA substrates, or (3) t- elements with true CCA tails are aminoacylated by aaRSs, inhibiting their detection with RNA sequencing because ligation is blocked by the resulting amino acid at the 3′ end. A further uncertainty about t-element metabolism is whether they are recognized by the enzymes responsible for post-transcriptional base modifications and/or aminoacylation of tRNAs. Therefore, a more comprehensive understanding of t-element processing and function necessitates a detailed investigation of their relationships with the full suite of tRNA-interacting enzymes.

Recent advances have been made in sequencing-based methods that can quantify expression and determine aminoacylation state for tRNAs in a gene-specific fashion (Evans et al. 2017; Pavlova et al. 2020; Behrens et al. 2021; Watkins et al. 2022; Davidsen and Sullivan 2024). These approaches incorporate a classic method to distinguish between aminoacylated and uncharged tRNAs that has been termed the “Whitfield reaction” (Davidsen and Sullivan 2024). This method involves treatment with sodium periodate to elicit ring opening and destabilization of the 3′ ribose followed by removal of the 3′ nucleotide by incubation under basic conditions. Because of the presence of an amino acid on the 3′ nucleotide of a tRNA protects the transcript from this reaction, there is a resulting length difference between aminoacylated tRNAs (full-length) and uncharged tRNAs (lacking their 3′ nucleotide).

In principle, the length difference generated by periodate treatment can be assessed on a global level with tRNA-seq. However, the extensive post-transcriptional base modifications in tRNAs (Cappannini et al. 2024) create substantial technical challenges because they can block reverse transcriptase (RT) progression during generation of cDNA (Wilusz 2015; Padhiar et al. 2024). Some tRNA-seq methods such as YAMAT-seq specifically capture full-length tRNAs by targeting transcripts with a 3′ NCCA overhang for double-stranded adapter ligation (Shigematsu et al. 2017). However, this approach produces a highly biased pool of sequences because it excludes tRNAs with base modifications that block RTs. The limitations of sequencing full-length tRNAs can be partially overcome through treatment with the dealkylating enzyme AlkB to remove some inhibitory base modification or use of specialized RTs with high processivity (Cozen et al. 2015; Zheng et al. 2015; Warren et al. 2021a; Scheepbouwer et al. 2023). However, these approaches have their own drawbacks because of the extensive RNA degradation introduced by AlkB buffer conditions and increased potential for RT artefacts. An alternative or complementary approach is to circularize cDNA or ligate a second adapter to its 3′ end after complementary DNA (cDNA) synthesis so that tRNA-derived sequences are captured regardless of whether they are full-length or truncated (Pinkard et al. 2020; Behrens et al. 2021; Watkins et al. 2022; Scheepbouwer et al. 2023).

In addition to the types of base modifications that inhibit RT processivity, tRNAs contain many modifications that lead to nucleotide misincorporations or indels by RTs during cDNA synthesis. The resulting sequence variants provide the opportunity to infer the location and identity of modified tRNA bases (Clark et al. 2016; Potapov et al. 2018; Enroth et al. 2019; Vandivier et al. 2019; Behrens et al. 2021; Warren et al. 2021a; Arrivé et al. 2023; Gołębiewska et al. 2024), and these modification patterns have been compared with aminoacylation levels to infer their role in tRNA charging (Hernandez-Alias et al. 2023).

Plants are fascinating models for studying tRNA metabolism because they maintain three parallel translation systems (nuclear/cytosolic, mitochondrial, and plastid). However, despite the advances in using tRNA-seq to measure aminoacylation levels, none of these approaches have yet been applied to plants. Here, we use periodate treatment in combination with a specialized tRNA-seq method (Watkins et al. 2022) to distinguish between aminoacylated and uncharged tRNAs and tRNA-like transcripts in the model angiosperm *Arabidopsis thaliana*. We also complement this analysis with previously published YAMAT-seq data (Warren et al. 2021a) to investigate the enigmatic plant mitochondrial t-elements.

## RESULTS

### MSR-seq captures all Arabidopsis tRNA isodecoder families

We used total RNA samples from *A. thaliana Col-0* rosettes (four weeks old, three biological replicates) to generate and sequence multiplex small RNA-seq (MSR-seq) libraries (Watkins et al. 2022) (Table S1). On average, 90.64% of the processed sequences mapped to a reference database consisting of annotated tRNAs in *A. thaliana* (Cognat et al. 2022), with 70.36%, 19.62%, and 0.66% of the processed sequences mapping to nuclear, plastid, and mitochondrial tRNA genes, respectively. All 220 unique tRNA gene sequences had mapped reads in one of the libraries, and all but two had mapped reads in all libraries (mapped read counts available via https://github.com/dbsloan/Arabidopsis_aminoacylation). Some of the reference sequences are similar enough that reads can produce equal mapping scores for two or more reference genes. When these ambiguously mapping reads are excluded, nine of the 220 reference genes lacked any coverage (eight tRNA-TyrGTA genes and one tRNA-SerAGA gene), but all isodecoder families (i.e., sets of tRNAs that share the same anticodon) found in *Arabidopsis* were still represented. Abundance estimates were highly correlated across biological replicates and different library treatment methods (Figure S1). Raw read abundances should be interpreted with caution because the tRNA sequencing process can result in heavily biased representation of specific genes, likely related to secondary structure, post-transcriptional base modifications, and other challenges associated with tRNA-seq (Ma et al. 2021; Warren et al. 2021a). In this dataset, tRNA-Trp is the most abundant isoacceptor family (i.e., a set of tRNAs charged with the same amino acid), accounting for an average of 19.6% of all nuclear tRNA reads. At the other extreme, tRNA-Cys (0.3%) and tRNA-His (0.8%) isoacceptors were very rare, accounting for less than 1% of all nuclear tRNA reads. Given the rarity of Trp in the proteome, it is unlikely that tRNA-Trp isoacceptors are as abundant as suggested by read counts or that the large differences among isoacceptors faithfully reflect biological abundances. Accordingly, relative comparisons across samples or treatment types may be the most informative interpretation of MSR-seq (and other tRNA-seq) datasets.

### CCA tail integrity and inferred aminoacylation levels vary among tRNAs and among genomic compartments

For MSR-seq library construction, we split the RNA samples from each biological replicate (see above) into three subsamples. One subsample was treated with periodate to distinguish between aminoacylated and uncharged tRNAs (see Introduction). The other two subsamples were used as a negative control (no periodate) or positive control (deacylation prior to periodate treatment) for the removal of the 3′ terminal nucleotide. Results from these three different treatments followed the expected pattern (Figure 1), as the overall percentage of mapped reads with an intact CCA tail in the test samples was intermediate (mean of 55.1%) between the negative control (75.7%) and the positive control (15.1%). Here and throughout, these “CCA percentages” were calculated after excluding reads lacking more than just a single 3′ nucleotide (see below for analysis of other read types that were missing additional 3′ sequence). The fact that the 3′ terminal nucleotide was not removed from 100% of tRNA reads in positive control samples indicates that the periodate treatment was not fully efficient in eliminating exposed 3′ nucleotides and/or that the preceding deacylation step was not fully efficient in removing the amino acids to expose these ends. Likewise, the fact that tRNA reads from negative control libraries did not exhibit 100% retention of an intact CCA tail indicates that a substantial percentage of transcripts in the *in vivo* tRNA pool lack the 3′ terminal nucleotide and/or that tRNAs were subject to some degradation during the RNA extraction and library construction process. These alternative interpretations are investigated below in the next two sections.

**Figure 1.**
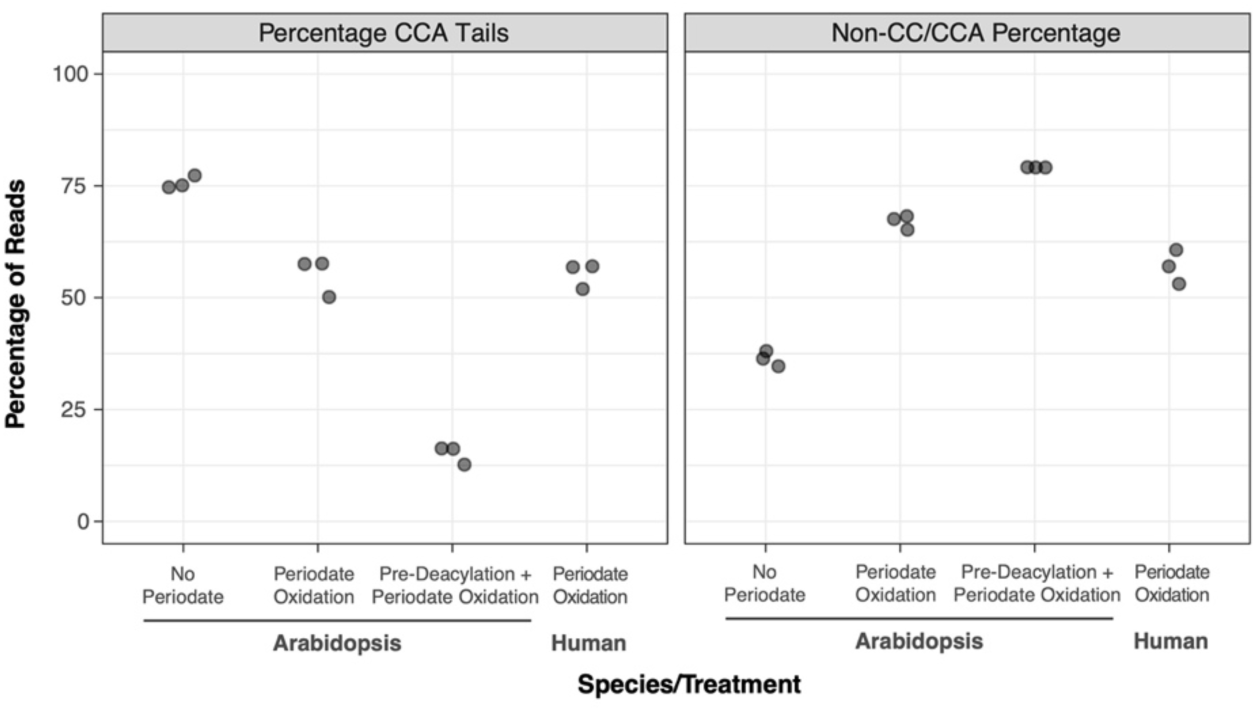
MSR-seq data from *Arabidopsis* (this study) and human cell line tRNAs (Watkins et al. 2022). Periodate treatment prior to MSR-seq library construction was used to estimate aminoacylation levels based on protection of the terminal nucleotide of the CCA by the amino acid. Reported values for CCA tails (left) represent the percentages of reads with intact CCA tails after excluding reads that lacked more than just a single 3′ nucleotide. In contrast, non-CC/CCA percentages (right) represent the fraction of *all* tRNA reads that lack two or more nucleotides at their 3′ ends, so values between the two panels do not sum to 100%. Such reads represent a large percentage of the dataset, especially in periodate-treated libraries. Three biological replicates were sequenced per treatment, with highly significant differences for both CCA tail (*p* = 6.3e-7, one-way ANOVA) and non-CC/CCA percentages (*p* = 5.1e-8, one-way ANOVA). The (periodate-treated) human samples are very similar to their *Arabidopsis* counterparts in terms of both CCA percentage and non-CC/CCA percentage.

In the periodate-treated test sample, inferred aminoacylation levels dipered substantially among tRNA genes. The percentage of reads with intact CCA tails ranged over two-fold across isoacceptor families (35.97% to 77.75%; Figure 2). Interestingly, the CCA percentage was significantly higher for tRNAs encoded in organellar genomes than in nuclear genomes when calculated as averages across isoacceptor families (Figure 2) or isodecoder families (Figure S2). Based on this observation, we reanalyzed a previously published MSR-seq dataset from human cell lines (Watkins et al. 2022) and found a similar pattern, with significantly higher percentages of intact CCA tails in mitochondrial tRNAs than in nuclear-encoded tRNAs (Figure S3).

**Figure 2.**
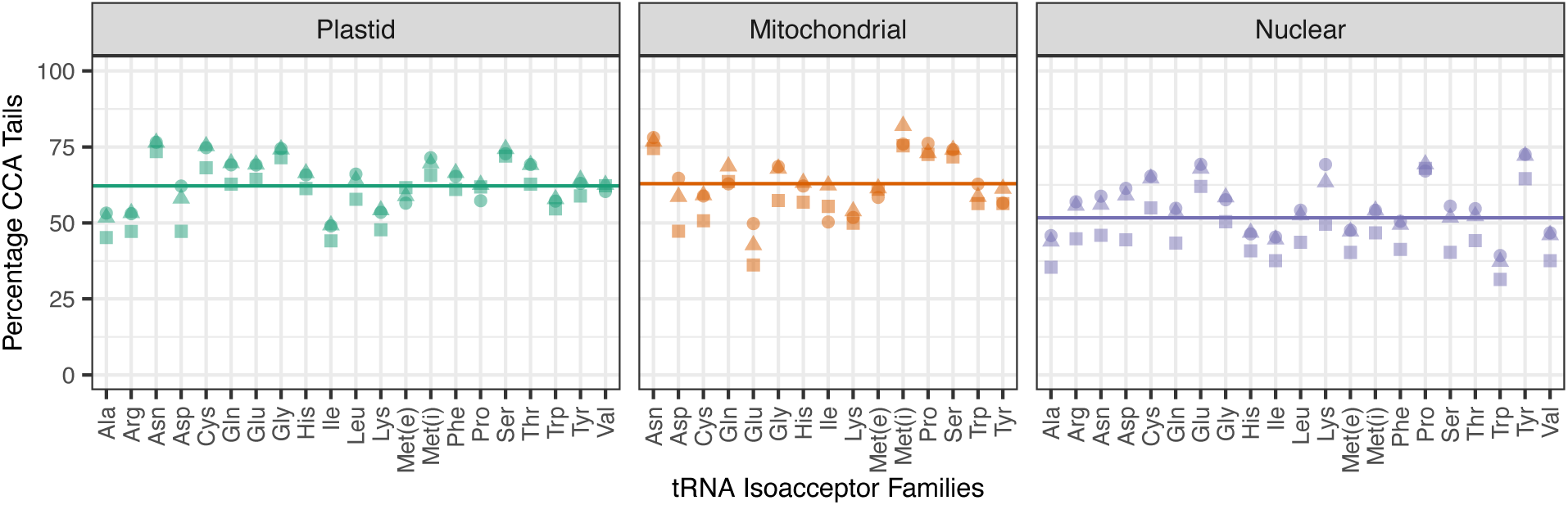
Variation among *Arabidopsis* tRNA isoacceptor families and genomic compartments in CCA tail integrity in response to periodate treatment. Reported values represent the percentages of reads with intact CCA tails after excluding reads that lacked more than just a single 3′ nucleotide. Biological replicates are indicated by different shapes. The average frequency of intact CCA tails differs significantly across genomic compartments (*p* = 0.0005; one-way ANOVA), with a lower rate for nuclear-encoded tRNAs than organellar tRNAs, as indicated by horizontal lines (means) in each panel. Met(e) and Met(i) refer to elongator and initiator tRNA-Met genes, respectively. We also found a similar difference between organellar and nuclear reads when retention of CCA tails was expressed as a percentage of total reads and not relative to just reads ending in CCA or CC (*p* = 1.5e-7; one-way ANOVA). A similar analysis subdivided into isodecoder families is presented in Figure S2.

The premise of using periodate treatment to infer aminoacylation levels is that uncharged tRNAs are susceptible to removal of their terminal 3′ nucleotide. However, we found that tRNAs in negative control (no periodate) libraries had frequencies of intact CCA tails that were strongly correlated with the periodate-treated test samples (*r* = 0.77; *p* = 8.9e-40), albeit with the control libraries at higher levels (Figure 3a). Accordingly, organellar tRNAs had a higher percentage of intact CCA tails than nuclear-encoded tRNAs even without periodate treatment (Figure 3a) (*p* = 6.9e-9; *t*-test). This observation implies that nuclear-encoded tRNAs have lower aminoacylation levels, in part because a smaller fraction of them within the cellular pool have an intact CCA tail. It is also possible that these tRNAs are more vulnerable to degradation during RNA extraction and library construction for some reason (including that they might be less likely to be protected by an amino acid at their 3′ end). Therefore, to further investigate variation in aminoacylation levels, we repeated the comparison of CCA tail percentage between nuclear and organellar tRNAs, after dividing the value for each gene under periodate treatment by the corresponding value from no- periodate controls. This analysis also found higher values for CCA tail frequencies for organellar tRNAs than for nuclear tRNAs (*p* = 1.3e-5; *t*-test), meaning that organellar tRNAs are more likely to keep their CCA tail even when adjusting for their higher starting percentage of CCA tails.

**Figure 3.**
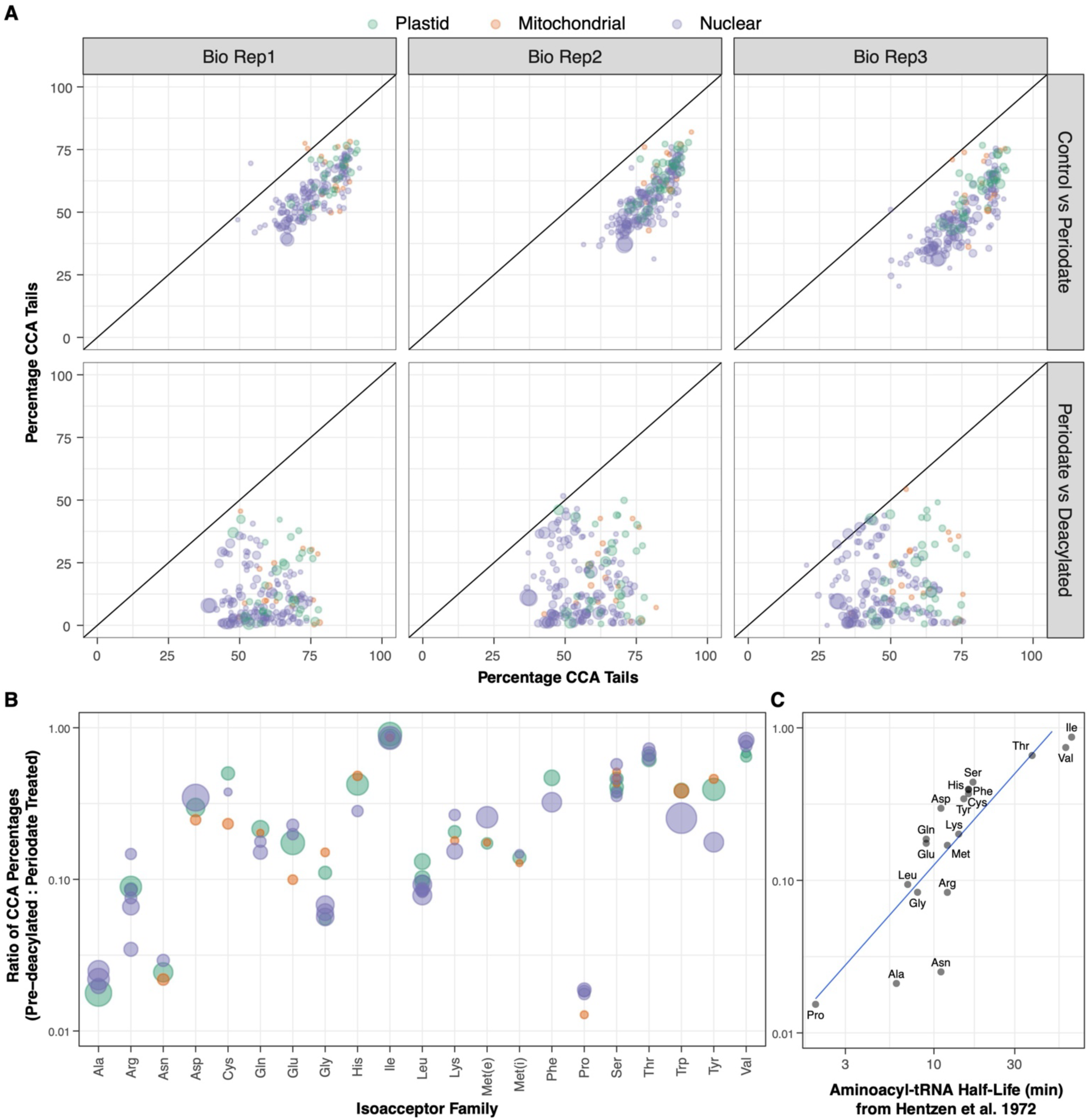
Comparison of CCA tail retention across treatments. (A) Correlation in percentage of *Arabidopsis* reads with intact CCA tails are shown for each biological replicate. Reported values represent the percentages of reads with intact CCA tails after excluding reads that lacked more than just a single 3′ nucleotide. The top row shows the comparison between the negative control libraries (no periodate) on the x-axis and the periodate-treated test samples on the y-axis. The bottom row shows the comparison between the periodate-treated test samples on the x-axis and the positive control samples (pre-deacylated prior to periodate treatment) on the y-axis. Each point represents an individual tRNA gene (minimum of 100 reads per gene), with color indicating genome of origin and size proportional to the average number of reads mapping to that gene. A one-to-one line is plotted in each panel. (B) Values on y-axis represent the ratio of CCA percentages for the pre-deacylated treatment vs. the standard periodate treatment by isoacceptor family. A value of 1 indicates that pre-deacylation did not result in any reduction of CCA tail percentage, whereas a value of 0 indicates that pre-deacylation completely eliminated CCA tails. Each point represents an isodecoder family averaged across genes with a minimum of 100 reads in that family and three biological replicates. Point size reflects average read abundance for that isodecoder family. Met(e) and Met(i) refer to elongator and initiator tRNA-Met genes, respectively. (C) Comparison of CCA retention rates (unweighted average of values in panel B) with aminoacyl-tRNA estimated for *E. coli* tRNAs at 37 °C, pH 8.6 (Hentzen et al. 1972). These half-life estimates are strongly correlated with the observed retention of CCA tails in *Arabidopsis* after a 30 min pre- deacylation at 37 °C, pH 9.0 followed by periodate treatment (*p* = 7.9e-6, *r* = 0.84, log-transformed).

In contrast to the strong correlation in the frequency of intact CCA tails between test samples and negative controls, there was little relationship between test samples and positive controls (pre-deacylated with high pH treatment prior to periodate treatment; Figure 3a; *r* = 0.04; *p* = 0.55). In principle, pre-deacylation should make tRNAs universally susceptible to the periodate. Many tRNA genes showed the expected pattern, with intact CCA tail percentages dropping to ∼0% in the pre-deacylated libraries. However, many other tRNAs exhibited little or no difference when a pre-deacylation treatment was applied, suggesting that effectiveness of deacylation may vary across tRNAs and that longer incubations may be necessary for complete deacylation (Davidsen and Sullivan 2024). Certain isoacceptor families such as Ile, Thr, and Val showed especially high retention of intact CCA tails despite pre-deacylation treatment (Figure 3b). Overall, the retention of CCA tails following pre-deacylation and periodate treatment exhibited a striking correlation (Figure 3c) with previous estimates of aminoacyl-tRNA half-life for *E. coli* tRNAs under similar deacylation conditions (Hentzen et al. 1972).

### RNA extraction method has minimal effect on CCA tail integrity, but periodate treatment causes extensive RNA degradation

Many *Arabidopsis* tRNA reads were missing more than just the terminal 3′ nucleotide. The frequency of such reads was higher in periodate-treated samples than in negative controls (no periodate) and increased further when we performed a deacylation step prior to treating with periodate (Figure 1). We also reanalyzed previously published MSR-seq data (Watkins et al. 2022) from human cell lines and found similar frequencies (Figure 1). The presence of shortened reads with incomplete CCA tails even in control libraries could reflect the true presence of these molecules *in vivo*, as CCA tail integrity can be responsive to environmental and subcellular conditions (Czech 2020; Lucas et al. 2023; Scheepbouwer et al. 2024). However, artefactual RNA damage may also contribute to this pool of shortened transcripts. Therefore, we performed a follow-up experiment to investigate the possible role of damage during RNA extraction and specifically the low pH in the acid-phenol extraction method that is typically used to preserve tRNA aminoacylation state (Zaborske et al. 2009). In this follow-up experiment, we generated libraries for *Arabidopsis* RNA samples that were extracted in parallel with either an acid-phenol protocol or a conventional Trizol protocol. We found that the results from acid-phenol and Trizol RNA samples were highly correlated (Figure S4) with significantly higher percentages of reads with intact CCA tails in the Trizol samples than in acid-phenol samples (*p* = 3.5e-32 and *p* = 2.3e-34 in control and periodate treatments, respectively). However, the difference between the extraction methods was very small (mean difference of 2.3% and 3.1% in control and periodate treatments, respectively). The two extraction methods did not differ in the percentage of shortened reads that lacked more than just the terminal 3′ nucleotide (*p* = 0.06 and *p* = 0.72 in control and periodate treatments, respectively). Therefore, the acid-phenol extraction approach does not appear to have caused a substantial increase in damage over conventional Trizol methods. However, somewhat surprisingly, the similar readouts from the different extraction methods and the fact that Trizol produced *higher* CCA tail percentages also suggest that our acid-phenol extractions may not be better than Trizol protocols at preserving the tRNA aminoacylation state.

To further investigate potential artefactual sources of RNA damage, we compared the integrity of total *Arabidopsis* RNA in a sample treated with a series of periodate, ribose, sodium tetraborate, and T4 polynucleotide kinase incubations according to the MSR-seq protocol (see Methods) to an untreated control sample. This comparison revealed extensive degradation in the treated sample (Figure S5), confirming that effects are not limited to removal of just a single 3′ nucleotide. Recent optimization efforts have shown that the desired outcome of periodate treatment can be obtained with much shorter periodate incubation times at lower temperatures and replacement of sodium tetraborate incubation with a less basic alternative to limit these additional forms of damage (Davidsen and Sullivan 2024).

### Periodate treatment results in near-complete removal of 3′ terminal nucleotide from synthetic spike-in control tRNAs

Given that deacylating RNA samples prior to periodate treatment did not lead to complete elimination of 3′ nucleotides (Figures 1 and 3), a more robust positive control for periodate-mediated removal would be desirable. Therefore, in the follow-up experiment that compared acid-phenol and Trizol extraction methods, we also included a synthetic “spike-in” tRNA that was added to each *Arabidopsis* RNA sample prior to library construction. Because these synthetic tRNAs are not aminoacylated, periodate treatment would be expected to fully remove terminal 3′ nucleotides. Indeed, reads derived from the spike-in exhibited near-complete elimination of the 3′ nucleotide, with an intact CCA tail percentage of only 1.7% in periodate-treated samples (Figure 4). This finding implies that the failure to completely remove 3′ nucleotides from some *Arabidopsis* tRNAs (Figures 1 and 3) was likely the result of incomplete deacylation rather than low efficiency of the periodate reaction itself.

**Figure 4.**
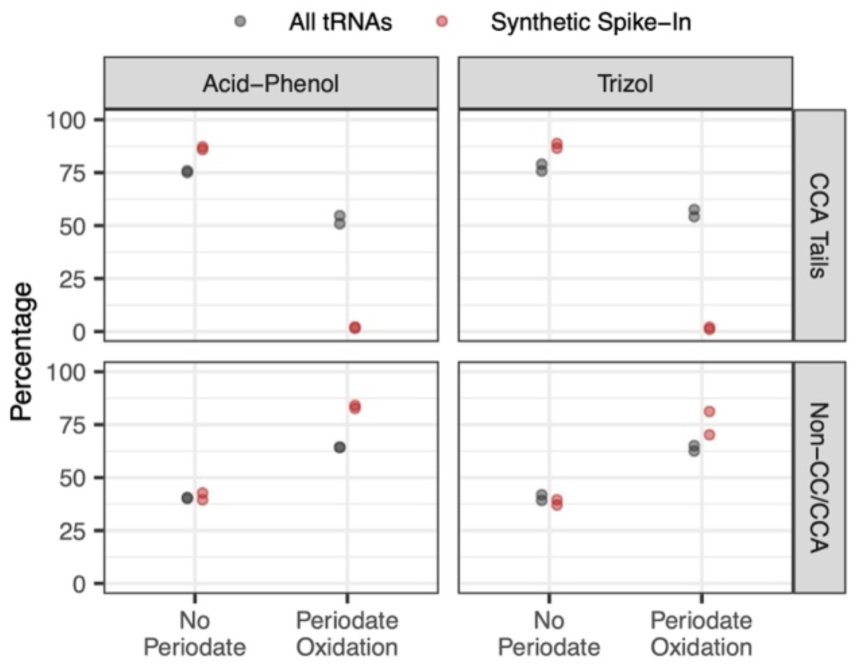
Inclusion of a synthetic tRNA spike-in control in MSR-seq libraries. Reported values for CCA tails (top row) represent the percentages of reads with intact CCA tails after excluding reads that lacked more than just a single 3′ nucleotide. In contrast, non-CC/CCA percentages (bottom row) represent the fraction of *all* tRNA reads that lack two or more nucleotides at their 3′ ends, so values between the panels do not sum to 100%. Values are shown for all *Arabidopsis* tRNAs (gray) and the synthetic spike-in tRNAs (red). Two biological replicates were performed for each of two RNA extraction methods: acid-phenol (left panels) and Trizol (right panels). For each biological replicate, control and periodate-treated libraries were generated.

The addition of a synthetic spike-in tRNA also provides insights into the origins of shortened reads that lack two or more nucleotides from their 3′ ends. In principle, the synthetic tRNAs should all retain intact CCA tails in control (no periodate) libraries, and they should only have a single 3′ nucleotide removed in periodate-treated libraries. We found that the CCA tail percentage in control libraries was higher for the synthetic tRNAs than for *Arabidopsis* tRNAs, but it still fell short of 100% (Figure 4). Furthermore, the majority of reads derived from the synthetic tRNAs in periodate-treated libraries lacked two or more 3′ nucleotides (Figure 4). This observation supports the conclusion that periodate treatment and downstream reactions can result in more extensive damage than the expected removal of a single 3′ nucleotide (Davidsen and Sullivan 2024). In addition, even in control libraries, there was a substantial frequency of these shortened reads derived from synthetic tRNAs (Figure 4). Thus, it appears that RNA handling and the process of library construction itself (e.g., overnight adapter ligation reaction) may result in some 3′ end degradation.

### Truncated tRNA sequences reveal ‘hard-stop’ base modification and tRNA fragmentation points

Some tRNA post-transcriptional base modifications inhibit RT progression (“hard stops”) and result in truncation of cDNA products at internal positions within the tRNA template (Wilusz 2015; Padhiar et al. 2024). In addition, there is an abundance of tRNA-derived fragments (tRFs) within cells (Keam and Hutvagner 2015; Lalande et al. 2020). Mapping the positions where sequencing reads start relative to the 5′ end of mature tRNAs can be used to infer positions of hard-stop base modifications and/or tRF breakpoints (Figures 5 and S6). We found that some *Arabidopsis* isodecoder families produced almost exclusively full-length tRNA sequences (e.g., nuclear tRNA-AspGTC and tRNA-GluTTC), whereas a few were almost always internally truncated (e.g., nuclear tRNA-LeuCAG and tRNA-LysTTT). Many others exhibited a mix of full-length and internally truncated reads (e.g., nuclear tRNA-AlaAGC and tRNA-SerGCA). Internal truncations were concentrated at the positions immediately 3′ of known base modifications (Figure 6). Because size selection in our library construction restricted the analysis to reads of at least ∼35 nt in length, truncations near the 3′ end of tRNAs were not investigated.

**Figure 5.**
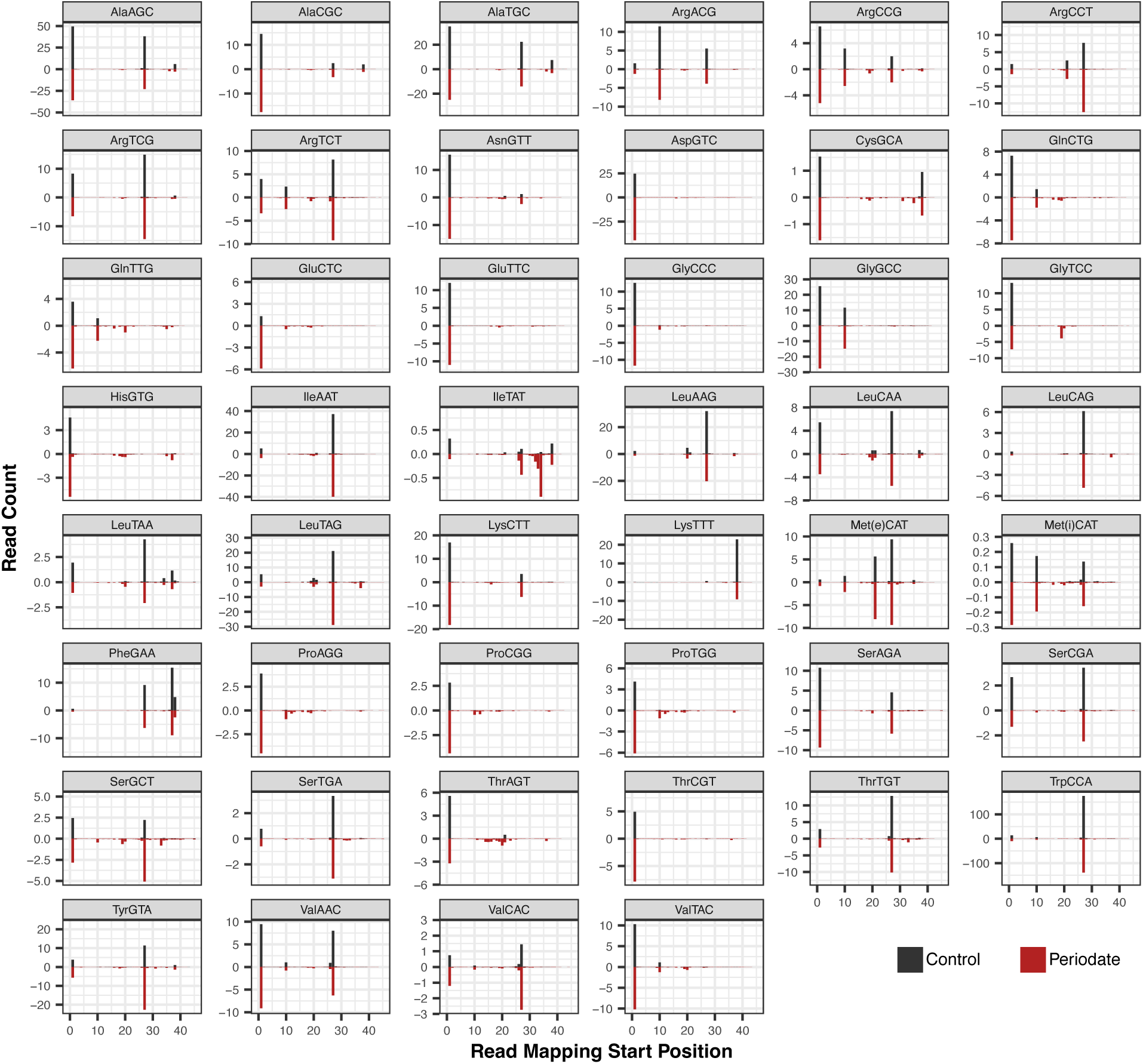
5′ mapping position of MSR-seq reads for all *Arabidopsis* tRNA isodecoder families found in the nuclear genome. Counts are represented per thousand reads that mapped to the nuclear tRNA gene set (averaged across the three biological replicates). Control libraries (black bars) are shown as positive values above the x-axis, while periodate-treated libraries (red bars) are shown as negative values below the y-axis. Mapping positions are standardized based on the Sprinzl coordinate system (Sprinzl et al. 1998).

**Figure 6.**
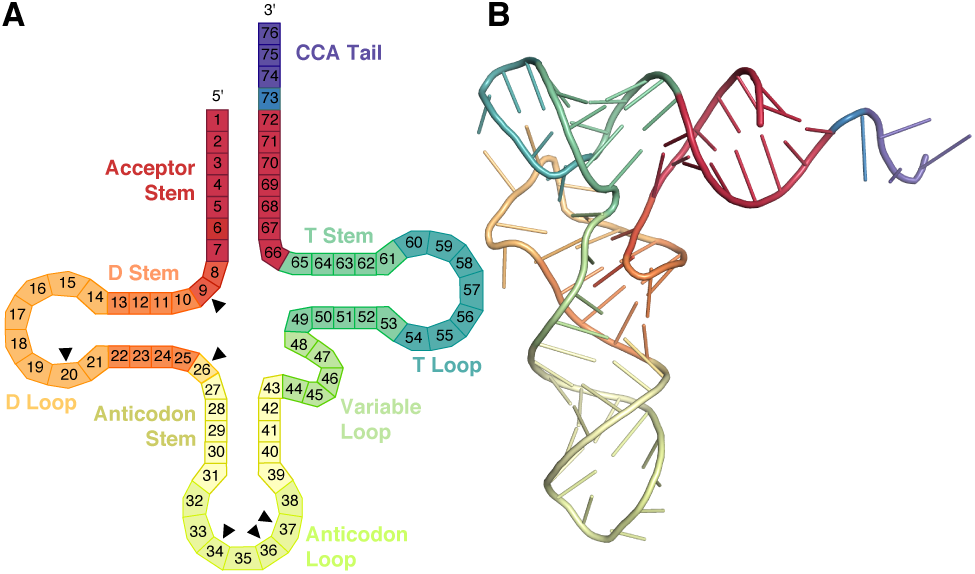
Canonical tRNA structures. (A) Secondary structure with bases numbered according to the Sprinzl coordinate system (Sprinzl et al. 1998). Black triangles indicate positions that were commonly associated with 5′ truncations. Note that because of size selection, short fragments resulting from truncations towards the end of molecule would not have been detectable. (B) Representative tRNA 3D structure (*Thermus thermophilus* tRNA-Glu; PDB 1G59), using the same color scheme as in panel A.

The distribution of 5′ mapping positions for a given isodecoder family was generally similar between periodate-treated and control libraries. However, nuclear tRNA-IleTAT (Figure 5) and plastid tRNA-LysTTT (Figure S6) exhibited intriguing exceptions. For these two isodecoder families, there was a major truncation point indicated by reads starting at nt position 34 (the wobble position of the anticodon) in periodate-treated samples, which was at very low frequency in no- periodate controls. We also found a large increase in nucleotide misincorporations or deletions at the neighboring position 33 in the subset of reads that are not truncated for these two isodecoder families in periodate-treated libraries (Figure S7). Similar observations in other species have been attributed to reactivity of periodate with 2-thio-modifications at the wobble position (Katanski et al. 2022; Davidsen and Sullivan 2024). More generally, many isodecoder families showed increases in the frequency of partial reads distributed across numerous low-frequency internal truncation points in periodate-treated libraries (Figures 5 and S6), suggesting more widespread effects of periodate treatment.

Mapping 3′ read end positions (Figures 7 and S8) can identify the breakpoints for 5′ tRFs (note that, unlike truncations at the 5′ end, shortening on the 3′ end of a read cannot be caused by hard-stop modifications and RT inhibition). Although most isodecoder families lacked major internal 3′ truncations points, there were some notable exceptions, such as nuclear tRNA-ValAAC (Figure 7) and plastid tRNA-IleCAT (Figure S8). When mapped at the level of individual genes, 5′ tRFs were most associated with breakpoints in the anticodon loops of nuclear tRNAs (Figure S9). However, it should be noted size selection during library construction limited detection of tRFs < ∼35 nt in length. Comparative mapping of end positions between periodate-treated and control libraries also illustrates the effect of periodate on 3′ nucleotide removal, as well as the sizeable population of reads that lack more than a single 3′ nucleotide (Figures 7 and S8).

**Figure 7.**
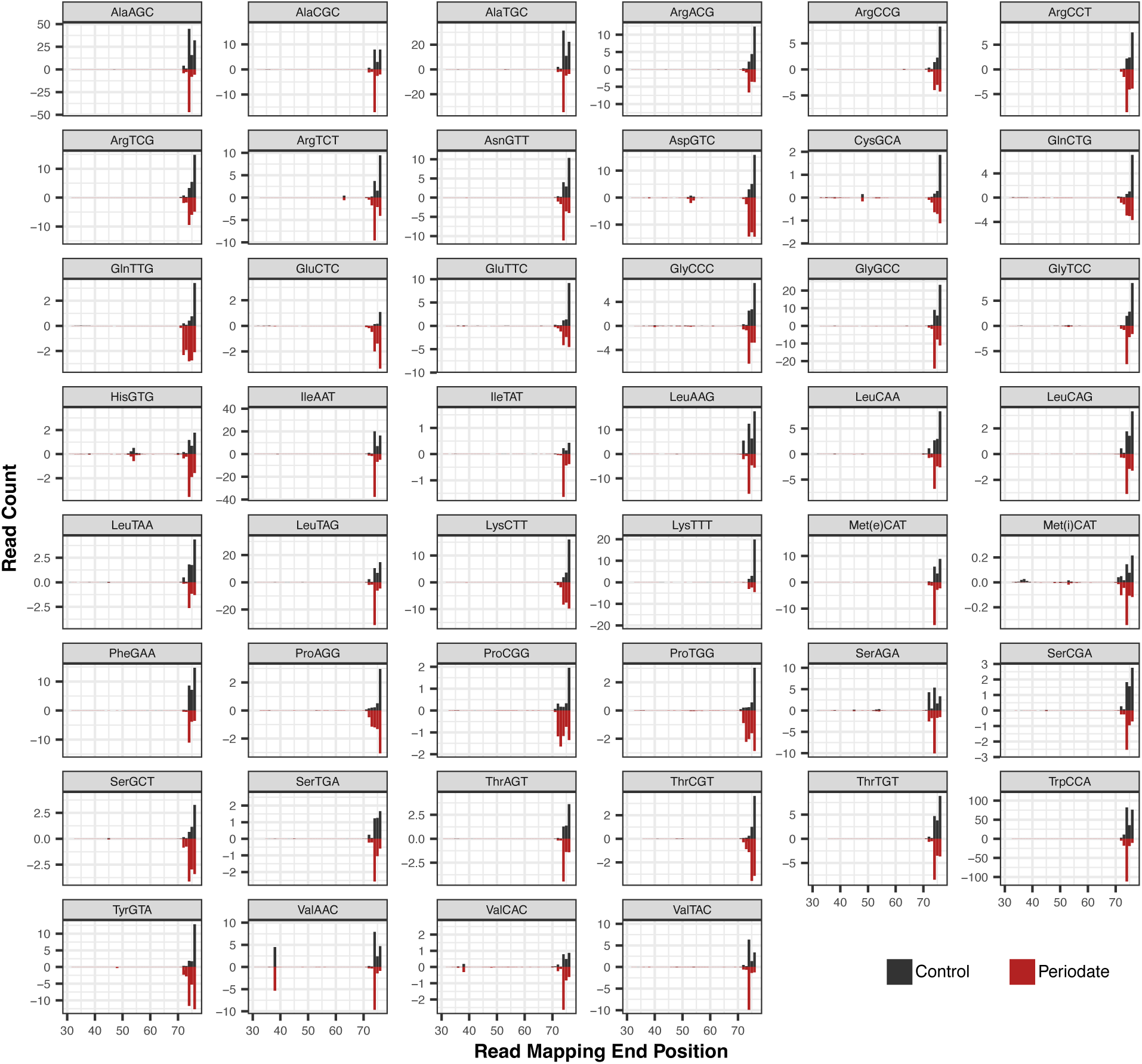
3′ mapping position of MSR-seq reads for all *Arabidopsis* tRNA isodecoder families found in the nuclear genome. Counts are represented per thousand reads that mapped to the nuclear tRNA gene set (averaged across the three biological replicates). Control libraries (black bars) are shown as positive values above the x-axis, while periodate-treated libraries (red bars) are shown as negative values below the y-axis. Mapping positions are standardized based on the Sprinzl coordinate system (Sprinzl et al. 1998).

### Post-transcriptional base modifications inferred from misincorporation patterns

Mapping the positions of nucleotide substitutions and indels onto reference sequences can identify bases that are post-transcriptionally modified (Clark et al. 2016; Potapov et al. 2018; Enroth et al. 2019; Vandivier et al. 2019; Behrens et al. 2021; Warren et al. 2021a; Arrivé et al. 2023; Gołębiewska et al. 2024). Using data from control (no periodate) libraries, we generated a global summary of position-specific misincorporation frequencies across all *Arabidopsis* tRNAs (Figure 8), revealing striking patterns such as the ubiquitous signature of the m^1^A58 modification in nuclear tRNAs (but not organellar tRNAs) and widespread misincorporations corresponding to the m^2^_2_G26 modification (Motorin and Helm 2010). Many other positions known to harbor modifications show high rates of misincorporations in multiple tRNAs (Suzuki 2021; Cappannini et al. 2024). It should be noted, however, that some modifications can be “silent” with regard to generation of RT misincorporations. Therefore, the absence of a misincorporation signature should not be taken as evidence that a base is completely free of modifications.

**Figure 8.**
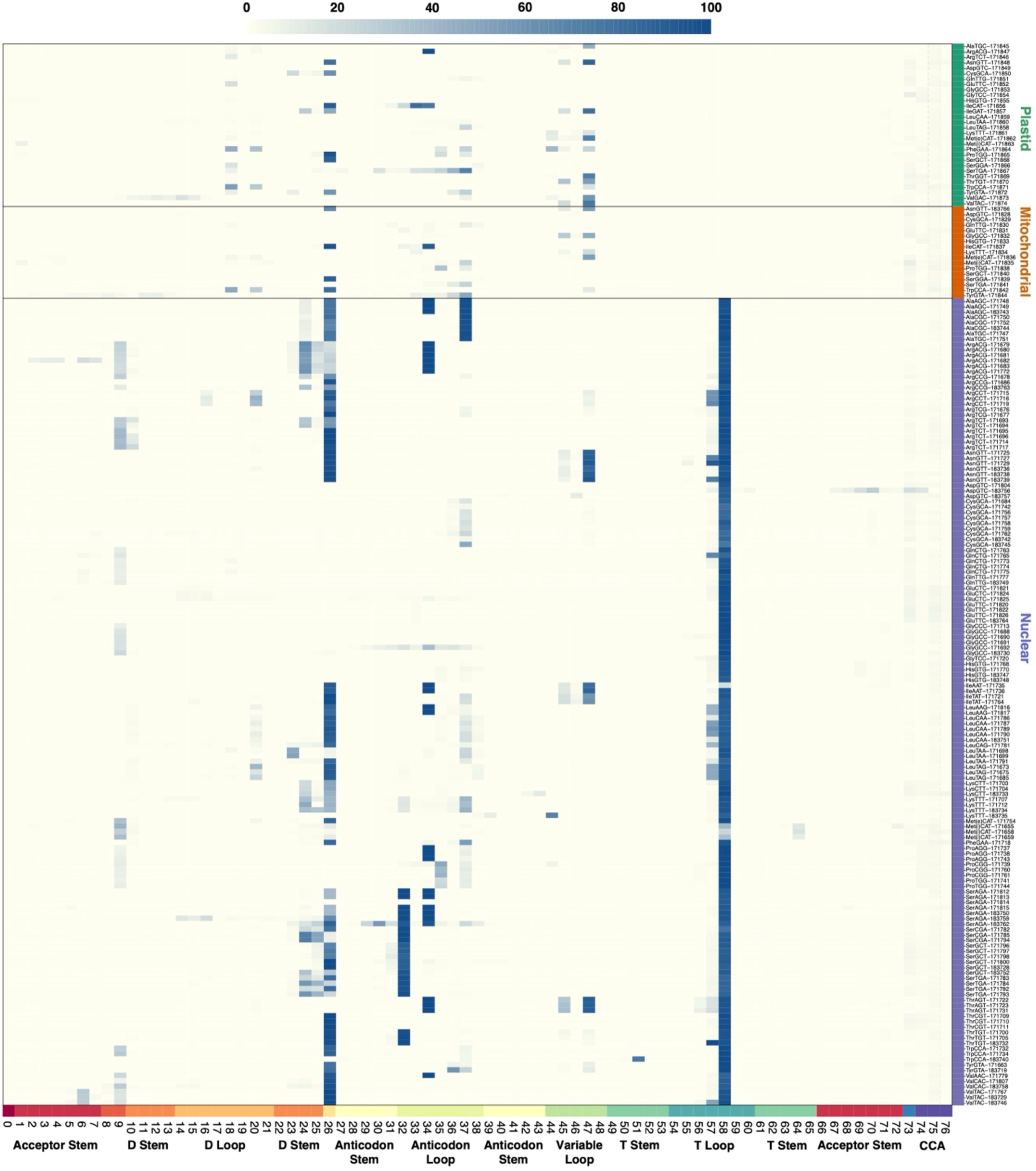
Heatmap representation of RT misincorporations across different tRNA genes and nucleotide positions in MSR-seq control (no periodate) libraries. The color scale reflects the average percentage of nucleotide substitutions and deletions for three biological replicates. Data are only reported for replicates with a read count >50. Mapping positions are standardized based on the Sprinzl coordinate system (Sprinzl et al. 1998), and corresponding structural positions are shown on the x-axis (see Figure 6 for definition of structural elements).

### Inferring relationships between base modification and aminoacylation rates

Base modifications can alter tRNA stability and molecular interactions. Therefore, they have the potential to affect aminoacylation rates (Hernandez-Alias et al. 2023). To investigate this possibility, we first tested for a difference in misincorporation rates in periodate-treated libraries between reads with intact CCA tails (aminoacylated tRNAs) and reads with CC tails (uncharged tRNAs). This analysis showed that misincorporation rates were highly correlated between these two read classes with no significant difference in rates between CCA and CC reads (*p* = 0.10; Figure S10). Therefore, misincorporation data did not provide evidence for an effect of single base modifications on aminoacylation rates.

We next considered 5′ truncations as another proxy for base modifications. In this case, we did find that reads with 5′ truncations had higher percentages of intact CCA tails than full-length reads in periodate-treated libraries (Figure S11). This pattern was observed for plastid (*p* = 2.5e- 11), mitochondrial (*p* = 1.1e-5), and nuclear (*p* = 2.4e-6) tRNAs, but the average difference in CCA percentages was approximately twice as large for organellar tRNAs than for nuclear tRNAs (Figure S11). The control (no periodate) libraries did not show this same pattern (*p* > 0.05 for each of the three genomes), which is consistent with the hypothesis that some of the base modifications that cause 5′ truncations during cDNA synthesis can facilitate aminoacylation. Thus, they run counter to the results from misincorporation data, which did not show an effect, and alternative interpretations should also be considered. For example, it is possible that the secondary structure of purified tRNAs is correlated with aminoacylation state and affects RT progression.

### Mitochondrial t-elements are transcribed, cleaved, base-modified, and CCA tailed – but not aminoacylated

The *Arabidopsis thaliana Col-0* mitochondrial reference genome (GenBank NC_037304.1) has eight annotated t-elements. To detect additional CCA-tailed transcripts that may be expressed and processed by tRNA-interacting enzymes, we reanalyzed a previously published *Arabidopsis* YAMAT-seq dataset (Warren et al. 2021a). This analysis detected expression and CCA-tailing of a family of tRNA-PheGAA-like sequences. One of these sequences is included in the plantRNA database (Cognat et al. 2022) as tRNA-TyrGTA-171843 because of a nucleotide substitution converting the GAA anticodon (Phe) to GTA (Tyr). However, this and other tRNA-Phe-like sequences in the *Arabidopsis* mitochondrial genome have been previously annotated as potential pseudogenes due to their divergence in sequence (Sloan et al. 2018). In total, there are seven unique tRNA-Phe-like sequences in the *Arabidopsis* mitochondrial genome (corresponding to 11 different loci due to additional identical duplicates). Some of these retain the GAA anticodon, whereas others share the derived GTA anticodon. We excluded one of these unique sequences from subsequent analysis because it represented only a small fragment and was not detected by YAMAT-seq. Analysis of YAMAT-seq data also found expression of a sequence that is 3′ of the mitochondrial *orf315*. This 63-nt transcript does not exhibit obvious homology to any tRNA gene, but it has sequence complementarity between its 5′ and 3′ ends, suggesting a “stem-loop” structure with a single-nucleotide 3′ overhang and added CCA tail. The same sequence is also present in the mitochondrial genomes of many other angiosperms.

Having cataloged the set of CCA-tailed mitochondrial transcripts, we then determined whether these were captured in our MSR-seq dataset. Overall, we found MSR-seq reads for five of eight previously annotated t-elements, all six of the analyzed tRNA-Phe-like sequences, and the *orf315*-adjacent sequence. In some cases, read abundance was extremely low, but we detected dozens to hundreds of reads per library for the 5′ *cox1* t-element, 3′ *ccmC* t-element, tRNA-Phe-like sequences, and the *orf315*-adjacent stem-loop (Dataset S1), enabling further investigation.

MSR-seq control (no periodate) libraries identified a diversity of post-transcriptionally added 3′ tails on the *ccmC* and *cox1* t-elements (Table 1) (Forner et al. 2007). The most common addition for the *ccmC* t-element was a canonical CCA tail, but many non-canonical variants were also detected. For the *cox1* t-element, a CCA tail was also abundant but slightly less common than a non-canonical CA tail. For reasons that are not clear, the distribution of post-transcriptionally added tails for the *cox1* t-element was notably different than observed in a previous analysis (Table 1) (Forner et al. 2007). Specifically, past sequencing of 19 cDNA clones did not identify any with a canonical CCA tail in contrast to the abundance of CCA tails in our dataset.

**Table 1.** Various post-transcriptional additions to the 3′ end of sequenced mitochondrial t-elements

| **3' Addition** | **Raw read counts in control (no periodate) libraries** |  |  |
| --- | --- | --- | --- |
|  | *ccmC* t-element | *cox1* t-element<br>(this study) | *cox1* t-element<br>(Forner et al. 2007) |
| CCA | 86 | 151 | 0 |
| CC | 35 | 47 | 3 |
| C | 12 | 19 | 4 |
| None | 0 | 12 | 6 |
| CA | 0 | 180 | 1 |
| CCAAA | 0 | 1 | 1 |
| CAAAA | 0 | 0 | 2 |
| CCAA | 4 | 4 | 0 |
| CCAC | 0 | 3 | 0 |
| CAAA | 0 | 1 | 0 |
| CAA | 0 | 3 | 0 |
| CACA | 0 | 1 | 0 |
| AC | 0 | 1 | 0 |
| A | 0 | 0 | 2 |

We also identified clear signatures of RT misincorporations in the *ccmC* t-element at the 26G nucleotide, a major target for post-transcriptional base modifications in tRNAs (Figure 9b). The mitochondrial *ccmC* t-element is homologous to plastid tRNA-IleCAT. The misincorporation profile at G26 is very similar between these two transcripts (dominated by deletions and G→C substitutions; Figure 9b). However, plastid tRNA-IleCAT also shows substantial hard-stop activity at this site, with a truncation rate of >50% at the 3′ nucleotide (position 27), whereas the *ccmC* t-element does not show any evidence of RT inhibition at the homologous position (Figure 9).

**Figure 9.**
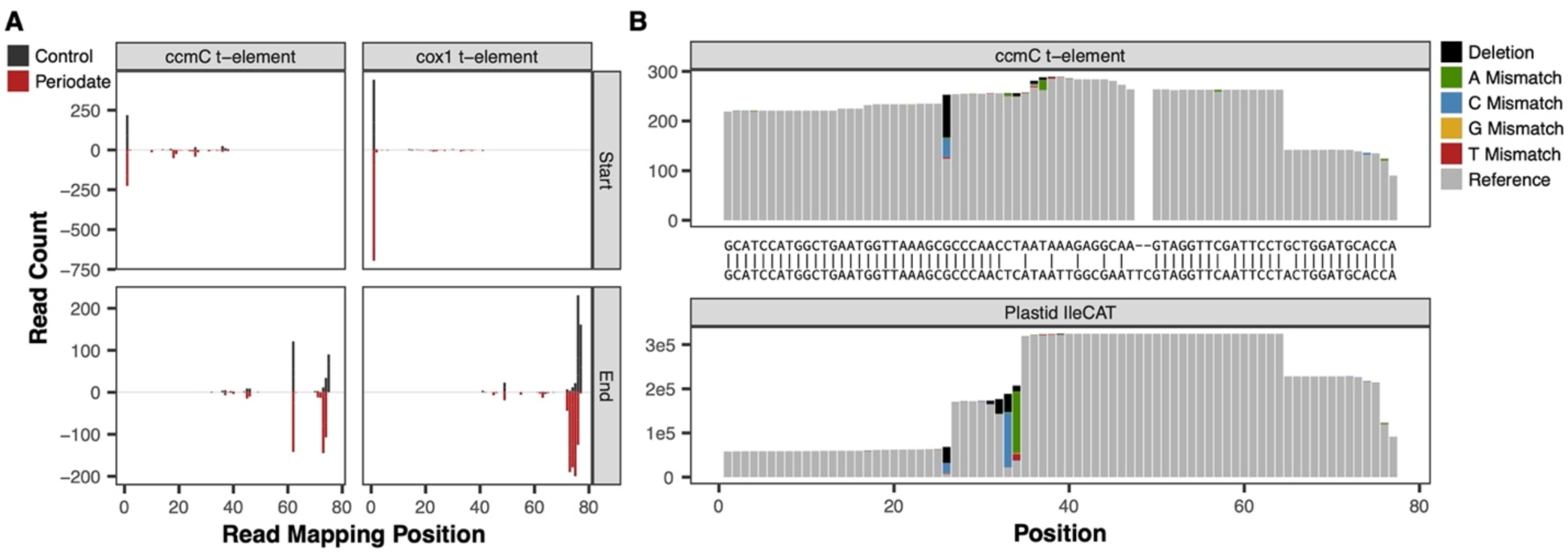
*Arabidopsis* mitochondrial t-element end-positions and misincorporations. The 5′ *cox1* and 3′ *ccmC* t- elements were both detected at substantial abundance in MSR-seq dataset. (A) The 5′ end (top) and 3′ end (bottom) mapping positions of reads are depicted in the same fashion as in Figures 5 and 7, respectively. Both *ccmC* (left) and *cox1* (right) t-elements are predominantly full-length on the 5′ end, whereas *ccmC* shows a high-frequency internal breakpoint among the 3′ ends. Periodate treatment leads to almost complete removal of the terminal 3′ nucleotide, indicating that the t-elements are not aminoacylated. (B) The mitochondrial *ccmC* t-element (top) is homologous to plastid tRNA-IleCAT (bottom). Nucleotide misincorporation patterns reveal an abundance deleted bases and nucleotide substitutions at G26 in both transcripts, indicating that the t-element likely shares the typical base modification at this position, although it appears to act as a “hard-stop” in tRNA-IleCAT but not in the t-element. We did not detect modified bases in the *cox1* t-element, which is not shown in panel B.

Therefore, it is not clear whether the *ccmC* t-element has the typical m^2^_2_G26 modification, although it is known that RT misincorporation and truncation patterns can vary depending on sequence context (Hauenschild et al. 2015; Behrens et al. 2021). The two homologs do share a high frequency of 3′ truncations after the 64U nucleotide (Figure 9). However, the t-element lacks the strong modification signatures that are present 33U and 34C in plastid tRNA-IleCAT, but that is not surprising because of divergence in the anticodon loop that distinguishes the two sequences (Figure 9b).

Overall, these observations provide evidence that mitochondrial t-elements and other tRNA-like sequences are recognized and processed by RNase P, RNase Z, CCAse, and enzymes that install post-transcriptional base modifications. However, our MSR-seq data indicate that the captured t-elements, tRNA-Phe-like sequences, and *orf315*-adjacent stem-loop were not aminoacylated because periodate treatment resulted in essentially complete elimination of the terminal 3′ nucleotide (Figure 9a; Dataset S1).

## DISCUSSION

### The effectiveness of MSR-seq in measuring plant tRNA expression and aminoacylation

The history of tRNA sequencing has long been a glass half-full, half-empty affair. On one hand, tRNAs played a pioneering role in the sequencing era, as a yeast tRNA was the first nucleic acid ever sequenced (Holley et al. 1965). On the other, they have arguably become the most challenging class of RNAs to accurately and quantitatively sequence. Recent innovations have begun to overcome this challenge, and our findings show that the MSR-seq workflow (Watkins et al. 2022) has promise for a more complete representation of plant tRNA pools. Because it can capture truncated RT products, this sequencing approach does not discriminate against transcripts carrying hard-stop base modifications. Accordingly, many tRNAs were found almost exclusively as truncated products in our dataset (Figures 5 and S6). These include isodecoder families that were essentially undetectable in a previous YAMAT-seq analysis of *Arabidopsis* (e.g., mitochondrial TyrGTA, SerTGA in all three genomes, and all nuclear Leu isodecoder families) and many others that were only detectable at appreciable levels with YAMAT-seq if samples were pre-treated with dealkylating enzyme AlkB to remove inhibitory base modifications (e.g., nuclear IleAAT, LysTTT, eMetCAT, PheGAA, ThrTGT, and TrpCCA) (Warren et al. 2021a).

Therefore, capturing truncated products resulting from RT inhibition greatly expands accessibility of the tRNA pool. However, there are obvious drawbacks to sequencing these truncated fragments. For example, unsequenced regions cannot be assessed for base modification profiles (Figure 8), and shorter sequences can be difficult to map unambiguously when multiple genes share similarity at their 3′ ends. In this study, our library size selection restricted the analysis to sequences of > ∼35 nt in length, so we did not investigate truncation patterns for extremely short fragments. Although treating RNA samples with the AlkB has been partially successful in circumventing inhibitory base modifications (Cozen et al. 2015; Zheng et al. 2015; Warren et al. 2021a; Scheepbouwer et al. 2023), the RNA degradation caused by its iron- containing buffer is likely to be especially problematic in the context of assays designed to use CCA-tail integrity as a proxy for aminoacylation state. Continued optimization of RT activity may instead be key to improving representation of full-length tRNAs. Recent strides have also been made to circumvent RT-associated challenges entirely by direct sequencing of tRNAs with nanopore or mass spectrometry technology (Thomas et al. 2021; Lucas et al. 2023; Sun et al. 2023; White et al. 2024; Yuan et al. 2024).

Our main objective in this study was to infer the plant tRNA aminoacylation landscape, and our results suggest that MSR-seq coupled with periodate treatment is a promising approach. In particular, this method appears to be highly effective at distinguishing a synthetic spike-in tRNA with no amino acid from functional biological tRNAs (Figure 4), which lends confidence to biological conclusions about tRNA-like sequences that do not appear to be aminoacylated and are, thus, unlikely to participate in translation (see below). However, our study also identified limitations associated with using periodate-induced read-length variation as a proxy for aminoacylation. The fact that current treatment conditions do not consistently remove one (and only one) nucleotide from uncharged tRNAs or consistently leave aminoacylated tRNAs intact makes the raw estimates of aminoacylation rates unreliable. This inconsistency also confounds the protective effect from amino acids with any true biological differences in tRNA end length.

Fortunately, an extensive effort was recently made to optimize periodate treatments for tRNA-seq (Davidsen and Sullivan 2024), which identified multiple factors that would likely reduce damage- related artefacts observed in our study. Potential modifications to our protocol include performing periodate treatment in the dark, reducing the periodate incubation temperature and duration, and using a lysine-containing buffer (pH 8.0) as a source of primary amine instead of the more alkaline sodium tetraborate treatment. Our data also suggest that incomplete deacylation can be a significant source of bias in tRNA-seq more generally (Figure 3), so optimization to ensure complete deacylation may also be important.

### Variation in aminoacylation rates between nuclear and organellar tRNAs

One unexpected result from our analysis was the difference in average CCA tail integrity between nuclear and organellar tRNAs (Figures 2 and S2). This observation raises the question as to whether these two classes have true biological differences in the standing ratio of aminoacyolated vs. uncharged tRNAs or differences in susceptibility to degradation that affects CCA tail integrity. The notion that aminoacylation rates and/or CCA tail integrity could differ substantially across subcellular environments is plausible. For example, it was recently shown that mammals sequester tRNAs in extracellular vesicle that differ in CCA tail integrity from the rest of the cellular tRNA pool despite similar base-modification profiles (Scheepbouwer et al. 2024). However, the potential for methodological artefacts should also be considered. We found similar differences in CCA tail integrity between nuclear and mitochondrial tRNAs from human cell culture (Figure S3) in a previously published MSR-seq dataset generated with the same periodate treatment protocol (Watkins et al. 2022). However, other tRNA-seq analyses of aminoacylation levels in human cell lines either did not see this same pattern (Evans et al. 2017) or even showed the opposite trend with lower inferred aminoacylation levels for mitochondrial tRNAs (Davidsen and Sullivan 2024). Therefore, future investigations will be needed to pinpoint the cause of this variation across studies. Regardless, focusing on gene-level differences will also be important because it appears that variation among genes is larger than the average difference between organellar and nuclear tRNAs.

### Functionality of mitochondrial t-elements and other tRNA-like sequences

To illustrate the utility of tRNA-seq approaches, we used our data to interpret the expression and functional interactions of mitochondrial t-elements. These tRNA-like sequences are already known to be transcribed and cleaved as part of expression with adjoining mRNA sequences, in addition to evidence for post-transcriptional addition of nucleotides to their 3′ end (Hanic-Joyce et al. 1990; Forner et al. 2007). We found that a full CCA tail was the most or one of the most common post-transcriptional additions (none of the t-elements have a genomically encoded CCA tail) (Table 1).

This observation suggests that the 3′ additions are mediated by a conventional CCAse (von Braun et al. 2007) despite evidence for substantial non-canonical tailing (Forner et al. 2007). Notably, some of the annotated t-elements in the *Arabidopsis* mitochondrial genome were rarely or never detected by either YAMAT-seq or MSR-seq (Dataset S1), raising the possibilities that they are either expressed at very low levels (or not at all) or rapidly degraded.

In addition to CCA-tailing, our data show that mitochondrial t-elements can have post-transcriptional base modifications (Figure 9b), implying an even larger set of tRNA-interacting enzymes are capable of recognizing t-elements. Importantly, however, we see almost complete elimination of t-element terminal 3′ nucleotides in response to periodate treatment (Figure 9a; Dataset S1). Despite caveats about inferring aminoacylation levels (see above), the extreme difference in CCA tail integrity in periodate-treated libraries relative to actual tRNAs that are known to function in translation appears to be strong evidence that these transcripts are rarely, if ever, recognized and charged by aaRS. We also found that the expressed family of tRNA-Phe-like sequences and the *orf315*-adjacent stem-loop in the mitochondrial genome show little or no evidence of aminoacylation. It is, thus, unlikely that any of these elements play a canonical tRNA role as decoder molecules in translation, but whether they play some other mitochondrial or cellular function remains to be determined.

These findings highlight the level of caution and specificity that is needed in interpreting functionality of tRNA(-like) genes. Although it is tempting to use transcription, cleavage, CCA-tailing, and post-transcriptional modifications as evidence for a functional role in translation, it is important to test for aminoacylation and, even better, loading of tRNAs onto ribosomes (Chen and Tanaka 2018) to make such determinations. Fortunately, the rapid and ongoing advances in specialized tRNA-seq methods is creating opportunities for such comprehensive assessments of tRNA metabolism.

## METHODS

### Plant growth and RNA extractions

*Arabidopsis thaliana* Col-0 seeds were surface sterilized for 10 min in a 50% bleach solution with Tween 20 (VWR 0777A) followed by three washes with sterile dH_2_O. They were then sown onto MS-agar plates, consisting of 2.2 g of Murashige and Skoog basal salts, 0.5 g β-(N-morpholino)ethanesulfonic acid (MES), and 7.5 g of Phytoblend plant agar (PTP01, Caisson Labs) per liter. Plates were then sealed in parafilm and cold-stratified at 4 °C for two days before being transferred to an incubator for growth under a 16hr-8hr light/dark cycle. Plants were grown for 4 weeks for the first MSR-seq experiment. Another batch of seeds was sown and grown for 3 weeks for the second experiment. For each biological replicate (three in the first experiment and two for each of the two RNA extraction methods in the second experiment), one rosette was pinched off from its root system with forceps and flash frozen in liquid N_2_ (∼15 to 20 mg of tissue per rosette).

All samples in the first experiment and half of the samples in the second experiment were extracted with a modified version of an acid-phenol protocol (Zaborske et al. 2009). Flash-frozen tissue was immediately ground with a micropestle and resuspended in 500 μl of lysis buffer consisting of 300 mM NaOAc pH 4.5 (Thermo Fisher AAJ63669AE), 10 mM EDTA, pH 8.0 (Invitrogen AM9262), and 1% sodium dodecyl sulfate (SDS) in RNase-free dH_2_O. 500 μl of an acid-phenol, chloroform, isoamyl-alcohol mixture (125:24:1) pH 4.5 (Invitrogen AM9720) was then added and followed by three bouts of ∼15 to 30 sec of vortexing, keeping samples on ice between bouts. Samples were then centrifuged for 15 min at 20,000 rcf and 4 °C. The resulting aqueous was then transferred to an additional 500 μl of the acid-phenol, chloroform, isoamyl-alcohol mixture and vortexed, followed by centrifugation under the same settings as above. After centrifugation, the aqueous phase was collected and ethanol precipitated, and the final pellet was dissolved in 20 μl of 10 mM NaOAc pH 4.5. The other half of the samples from the second experiment were extracted with Trizol (Ambion 15596018), following manufacturer’s instruction. The final pellet from each Trizol extraction was dissolved in 20 μl RNase-free dH_2_O.

### MSR-seq library construction and sequencing

Sequencing libraries were generated following the published MSR-seq protocol (Watkins et al. 2022) with minor modifications. Specific differences relative to the published version included the following: (1) we added 0.1% Tween 20 (VWR 0777A) in the high-salt buffer, (2) we used a single capture hairpin oligo (CHO; Table S2), so each library was processed in a separate tube with no multiplexing until the conventional Illumina indexes were incorporated at the PCR stage, (3) prior to PCR amplification, beads with ligated adapter molecules were stored overnight at -20 <u>°</u>C because we found that doing so resulted in more consistent amplification across samples, and (4) after amplification, libraries were pooled for subsequent size selection on a BluePippin (Sage Science) with a 3% cassette and Q3 Marker, using “Range” settings and size limits of 170-275 bp. A detailed version of the protocol used in these experiments is available via GitHub (https://github.com/dbsloan/Arabidopsis_aminoacylation).

In our first experiment, RNA samples were subjected to one of three different treatments:

1. standard periodate treatment to infer aminoacylation state, (2) periodate treatment preceded by a deacylation treatment, and (3) a no-periodate control treatment. Deacylation reactions (which were required for treatments #2 and #3) involved a 30 min incubation at 37 °C in 50 mM Tris-HCl pH 9.0, followed by ethanol precipitation. Periodate treatments were performed as described in the original MSR-seq publication (Watkins et al. 2022), using 500 ng of input RNA (either untreated or pre-deacylated). For the no-periodate controls, 1 μg of deacylated RNA was used as input for the library construction protocol.

In our second experiment, only the standard periodate and no-periodate control treatments were performed (i.e., there was no pre-deacylation treatment group), using RNA samples from the two different extraction protocols (acid-phenol and Trizol). We also included an internal control in these libraries by spiking in 1 ng of a synthesized tRNA (with the sequence of *Bacillus subtilis* tRNA-Ile [GenBank NC_000964.3: 31932-32008]; Table S2). This spike-in was added immediately prior to the periodate or deacylation step in the respective treatment.

After amplification and size selection, the final library pools were sent to Novogene for 2×150 bp paired-end sequencing on an Illumina NovaSeq platform. Raw reads for each library have been deposited into the NCBI Sequence Read Archive (SRA) (Table S1).

### MSR-seq read processing and mapping

All scripts used in read processing and subsequent data analysis are available via our GitHub repository (https://github.com/dbsloan/Arabidopsis_aminoacylation). BBMerge (Bushnell et al. 2017) was used to combine each pair of R1 and R2 reads into a single sequence with the following parameters: ordered=t qtrim=r minoverlap=30 mismatches=4. Merged reads were then further trimmed with a custom script to remove the adapter sequence from the MSR-seq CHO. Read pairs that that could not be combined by BBMerge or did not have the expected CHO adapter sequence (Table S1) were excluded from downstream steps.

All successfully processed reads were mapped against a reference that consisted of the unique *Arabidopsis thaliana* Col-0 tRNA sequences in the plantRNA database (Cognat et al. 2022) as of December 20, 2023. Later versions of our reference set also included the synthetic spike-in *Bacillus subtilis* tRNA-Ile sequence and mitochondrial t-element and stem-loop sequences (see below). Mapping was conducted with Bowtie v2.2.5 (Langmead and Salzberg 2012), using the following sensitive parameters to facilitate detection of reads with mismatches and indels that were introduced due to post-transcriptional base modifications: -p 12 -L 10 -i C,1 --mp 5,2 -- score-min L,-0.7,-0.7. However, even with these sensitive settings, mapping MSR-seq reads to tRNA references can be problematic because RT terminal transferase activity results in extra nucleotides being added past the 5′ end of the original RNA template and because reads that extend beyond the ends of reference sequences are heavily penalized under the Bowtie global alignment settings. Therefore, we used a custom local alignment pipeline based on NCBI BLASTN 2.14.1+ (Camacho et al. 2009) to pre-process reads and trim 5′ extensions prior to Bowtie mapping. A custom script was then used to parse the Sequence Alignment Map (SAM) file generated by Bowtie, quantifying the number of reads that mapped to each reference sequence and determining whether they contained a full CCA tail or were lacking one or more 3′ nucleotides. This script was run both with and without an option to exclude reads that map equally well to two or more reference sequences.

### Detecting RT misincorporations and truncations

SAM files were also parsed to identify reads with mismatches or small indels relative to the reference sequence, which can be caused by RT misincorporations in response to post-transcriptionally modified bases in the RNA template (Clark et al. 2016; Potapov et al. 2018; Enroth et al. 2019; Vandivier et al. 2019; Behrens et al. 2021; Warren et al. 2021a; Arrivé et al. 2023; Gołębiewska et al. 2024). For a global analysis of variants across all reference tRNAs (Figure 8), we used a previously published variant calling workflow (Edera and Sanchez-Puerta 2021). We also used Perbase v0.9.0 (https://github.com/sstadick/perbase) for analysis of sequence variants in some specific tRNAs of interest (Figures 9 and S7). In addition, the mapping positions of 5′ read ends were also parsed from SAM files to infer internal truncation points, which may result from base-modifications that block RT progression or from strand breaks (i.e., 3′ tRFs). Scripts used to perform SAM file parsing and data visualizations are all available via our GitHub repository (https://github.com/dbsloan/Arabidopsis_aminoacylation).

### Sprinzl coordinate system

Although tRNAs share a typical structural organization comprised of multiple stems and loops (Figure 7), the exact nucleotide positioning is not identical across tRNAs because of differences in the lengths of some of these structural motifs and certain “variable” elements. Therefore, we applied a standardized numbering scheme (Sprinzl et al. 1998) that anchored gene-specific nucleotide positions to their position in the canonical tRNA structure, using the classification of structural elements from the plantRNA database (Cognat et al. 2022) and a custom script (https://github.com/dbsloan/Arabidopsis_aminoacylation).

### Analysis of mitochondrial t-elements, pseudogenes, and stem-loops

To investigate whether mitochondrial t-elements and other tRNA-like sequences are expressed, post-transcriptionally processed, and aminoacylated, we first used existing YAMAT-seq data (Warren et al. 2021a) with a workflow to identify and quantify mitochondrially mapping reads that do not overlap with annotated tRNAs or other genes as previously described (Warren et al. 2021b). Based on the results of this analysis, we included the eight annotated t-elements in the *Arabidopsis* reference mitochondrial genome (NC_037304.1), as well as a set of tRNA-Phe-like sequences, and an expressed stem-loop associated with *orf315* in our tRNA reference set used for mapping (Dataset S1). This reference set is available via GitHub (https://github.com/dbsloan/Arabidopsis_aminoacylation). The MSR-seq mapping workflow described above was then re-run with this updated reference to quantify read abundance for these mitochondrial sequences and determine whether they retained intact CCA tails.

### Analysis of 5′ tRFs

To identify tRFs that represent the 5′ portion of tRNA genes, we parsed SAM files and screened for reads that terminated >7 nt from the annotated 3′ end of the reference tRNA sequence. We identified tRFs as high-frequency if the reads ending at that position represented >5% of all reads mapping to that tRNA. If other positions separated by three or fewer nt also cleared this threshold, we considered them to be a single tRF and assigned the tRF position to 3′- most site above this threshold. Customs scripts were used to apply these criteria (https://github.com/dbsloan/Arabidopsis_aminoacylation).

### Reanalysis of published MSR-seq data from human cell lines

To determine whether previously published MSR-seq data from human cell lines (Watkins et al. 2022) showed similar frequencies of non-CC/CCA reads and differences between nuclear and mitochondrial tRNAs in CCA percentages as we observed in *Arabidopsis*, we re-analyzed sequences obtained from NCBI SRA (SRR18302277, SRR18302278, and SRR18302279) with our pipeline. These three datasets were from control samples (i.e., no stress treatment), with a standard periodate protocol used in library construction (Watkins et al. 2022).

## DATA AVAILABILITY

MSR-seq reads have been deposited to NCBI SRA (BioProject PRJNA1154980). Custom scripts and processed data are available via GitHub (https://github.com/dbsloan/Arabidopsis_aminoacylation).

## ACKNOWLEDGEMENTS

We thank Lio Lee and Amanda Broz for assistance with plant growth and Chris Katanski for helpful methodological advice. This work was supported by funding from the National Science Foundation (MCB-2322154), an IUBMB Wood-Whelan Research Fellowship, and an HHMI Hanna H. Gray Fellowship.

---

**Figure S1.**
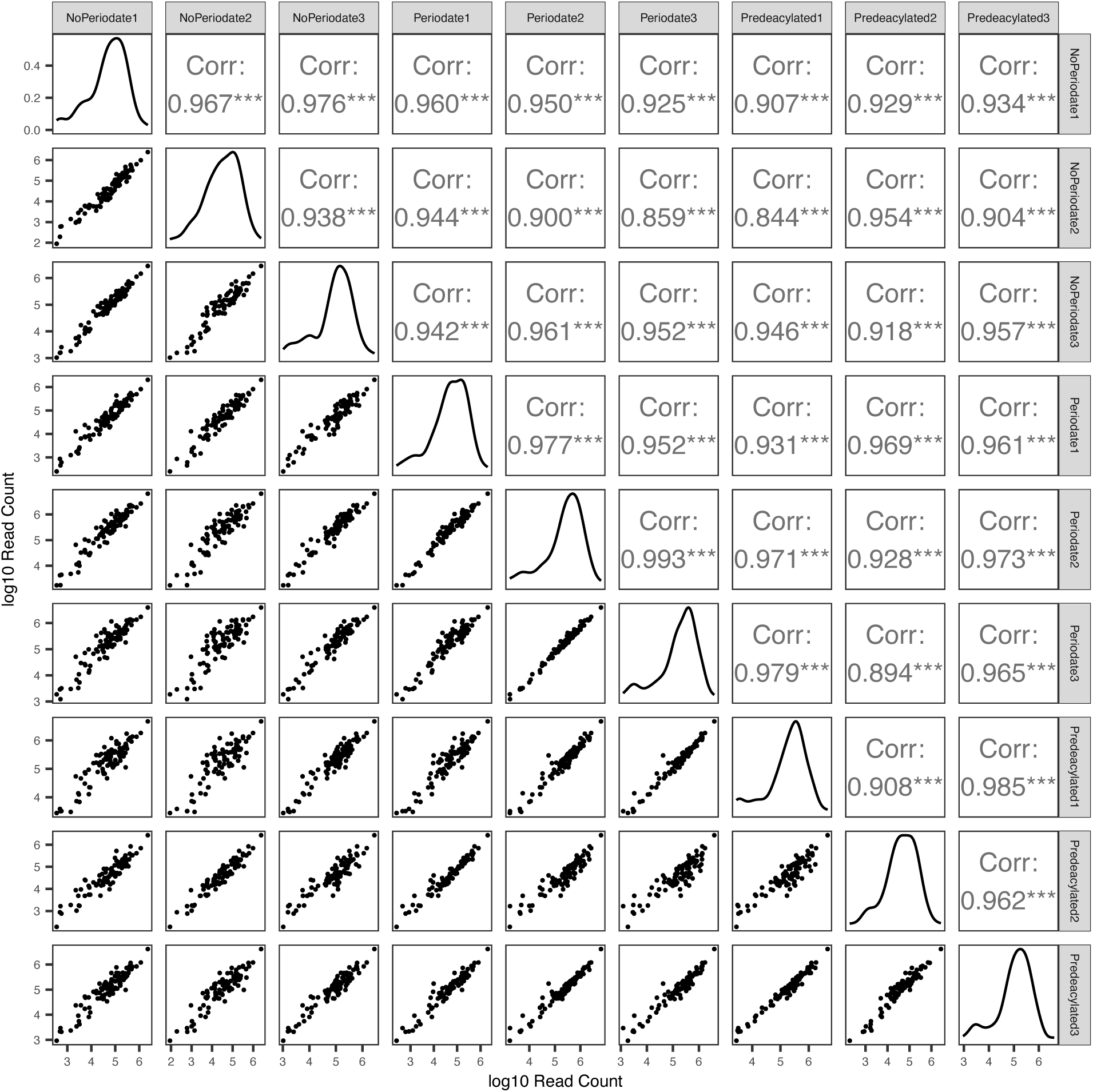
Pairwise matrix showing correlation in read abundance measurements for each library. In scatter plots below the primary diagonal, each point represents the summed read count for all members of an isodecoder family in a given genome (mitochondrial, plastid, or nuclear) on a log10 scale. Reported coefficients above the primary diagonal are from a Pearson correlation analysis on log10 abundance values. The plots on the primary diagonal itself report a read abundance density kernal for each library.

**Figure S2.**
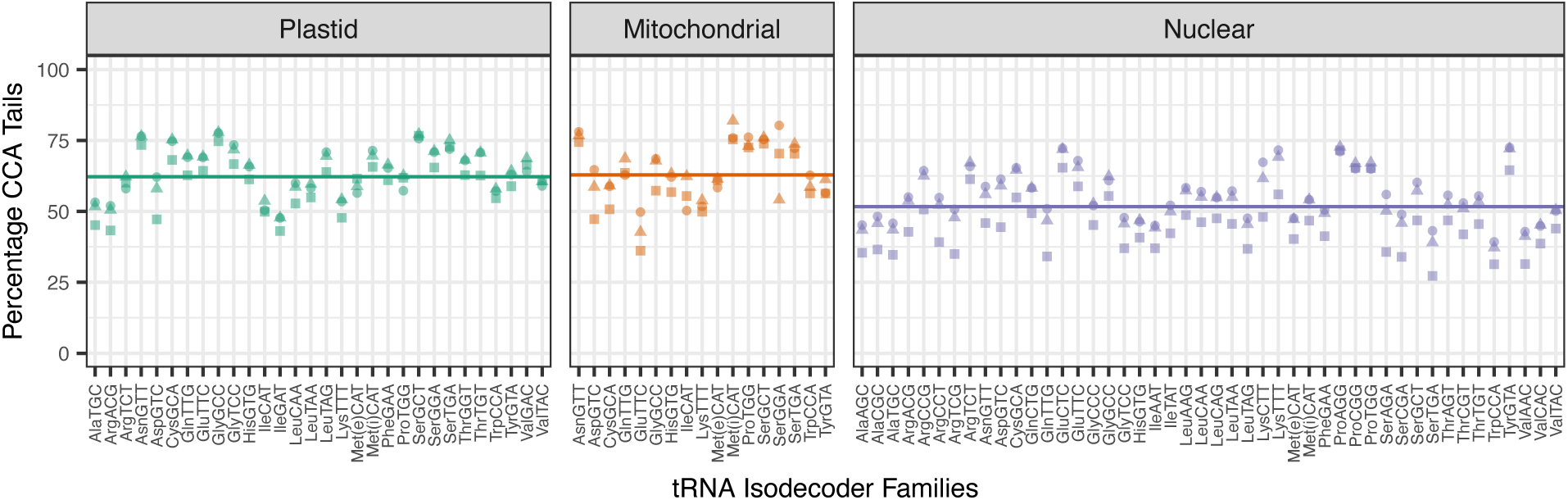
Variation among *Arabidopsis* tRNA isodecoder families and genomic compartments in CCA tail integrity in response to periodate treatment. Reported values represent the percentages of reads with intact CCA tails after excluding reads that lacked more than just a single 3′ nucleotide. Biological replicates are indicated by different shapes. The average frequency of intact CCA tails differs significantly across genomic compartments (*p* = 2.5e-7; one-way ANOVA), with a lower rate for nuclear-encoded than organellar tRNAs, as indicated by horizontal lines (means) in each panel. Met(e) and Met(i) refer to elongator and initiator tRNA-Met genes, respectively.

**Figure S3.**
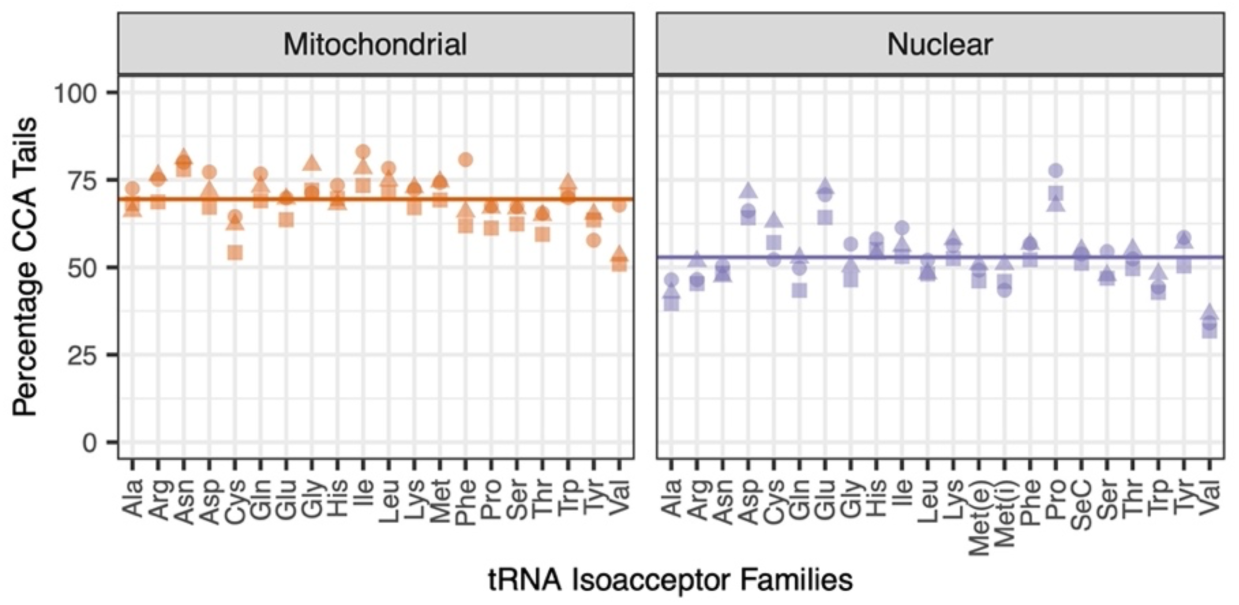
Variation among human cell line (HEK293T) tRNA isoacceptor families and genomic compartments in CCA tail integrity in response to periodate treatment. Data were reanalyzed from a previously published study (Watkins et al. 2022). Reported values represent the percentages of reads with intact CCA tails after excluding reads that lacked more than just a single 3′ nucleotide. Biological replicates are indicated by different shapes. The average frequency of intact CCA tails is significantly higher for mitochondrial tRNAs than nuclear-encoded tRNAs (*p* = 7.8e-9; *t*-test), as indicated by horizontal lines (means) in each panel. Met(e) and Met(i) refer to elongator and initiator tRNA-Met genes, respectively.

**Figure S4.**
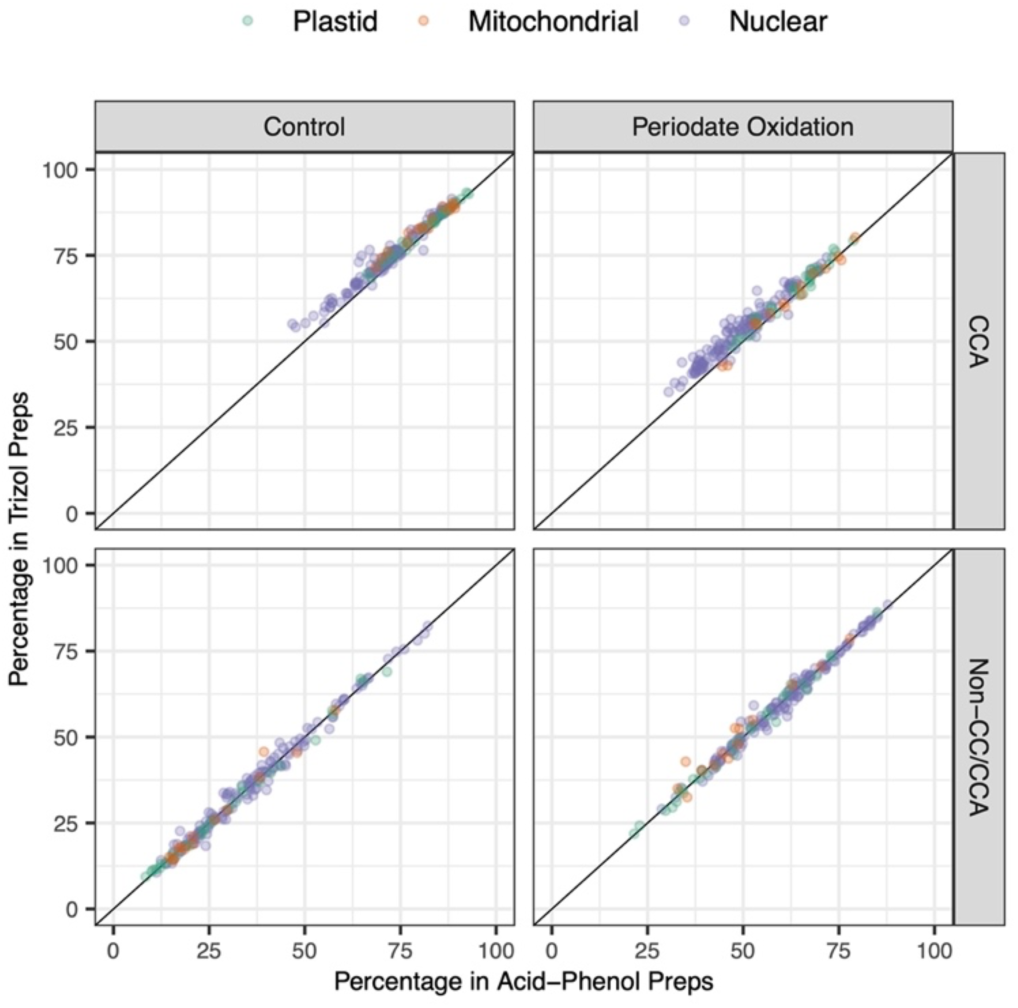
Comparison of acid-phenol and Trizol RNA extraction methods as input into *Arabidopsis* MSR-seq libraries. Reported values for CCA tails (top row) represent the percentages of reads with intact CCA tails after excluding reads that lacked more than just a single 3′ nucleotide. Non-CC/CCA percentages (bottom row) represent the fraction of all tRNA reads that lack two or more nucleotides at their 3′ ends. Each point represents an individual tRNA gene (minimum of 100 reads per gene), with color indicating genome of origin. A one-to-one line is plotted in each panel.

**Figure S5.**
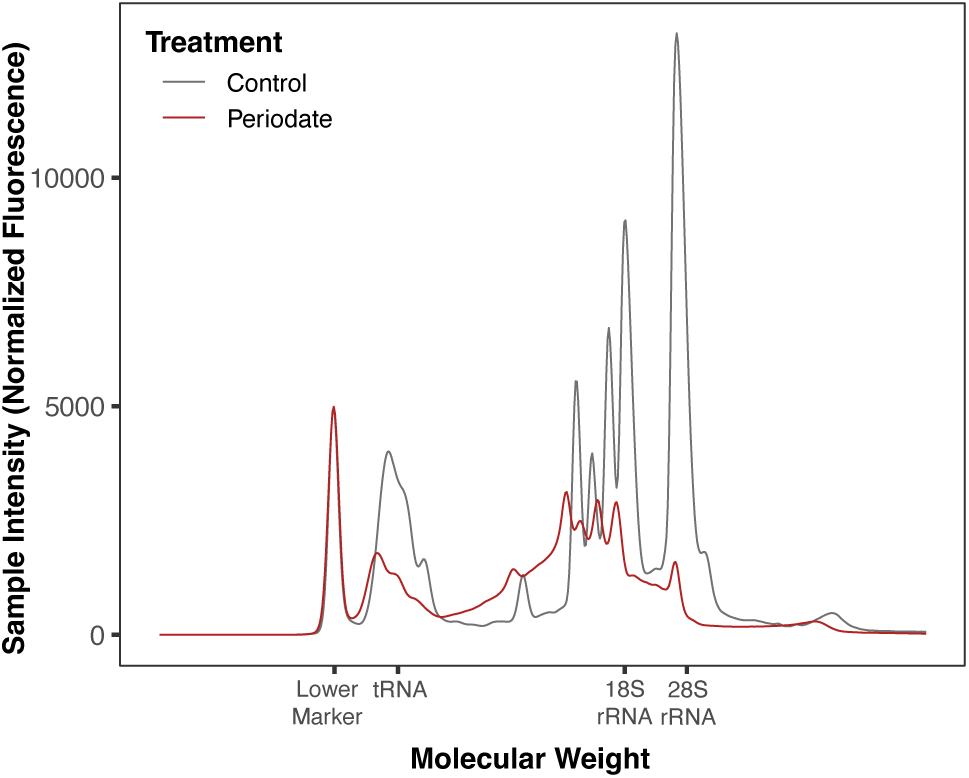
Comparison of RNA samples before (gray) and after (red) treatment with sodium periodate, ribose, sodium tetraborate, and T4 polynucleotide kinase according to the MSR-seq protocol. Data from an Agilent TapeStation 4150 show extensive RNA degradation from periodate treatment.

**Figure S6.**
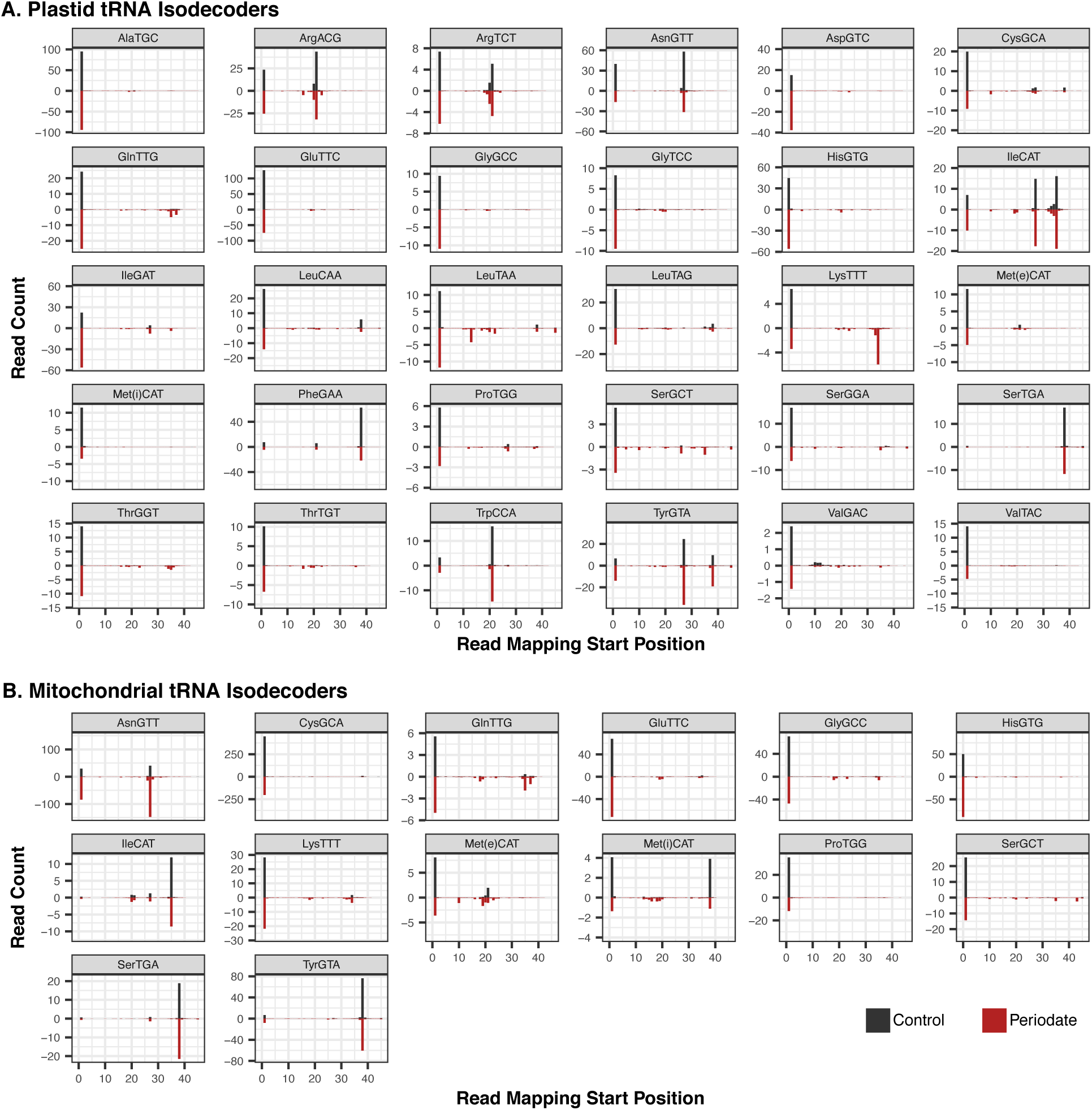
5′ mapping position of MSR-seq reads for *Arabidopsis* organellar tRNAs. Counts are represented per thousand reads that mapped to the (A) plastid or (B) mitochondrial tRNA gene set (averaged across the three biological replicates). Control libraries (black bars) are shown as positive values above the x-axis, while periodate-treated libraries (red bars) are shown as negative values below the y-axis. Mapping positions are standardized based on the Sprinzl coordinate system (Sprinzl et al. 1998). The mitochondrial AspGTC, SerGGA, and TrpCCA tRNAs are not included in this analysis because a large number of reads from their plastid-derived counterparts ambiguously map to these genes.

**Figure S7.**
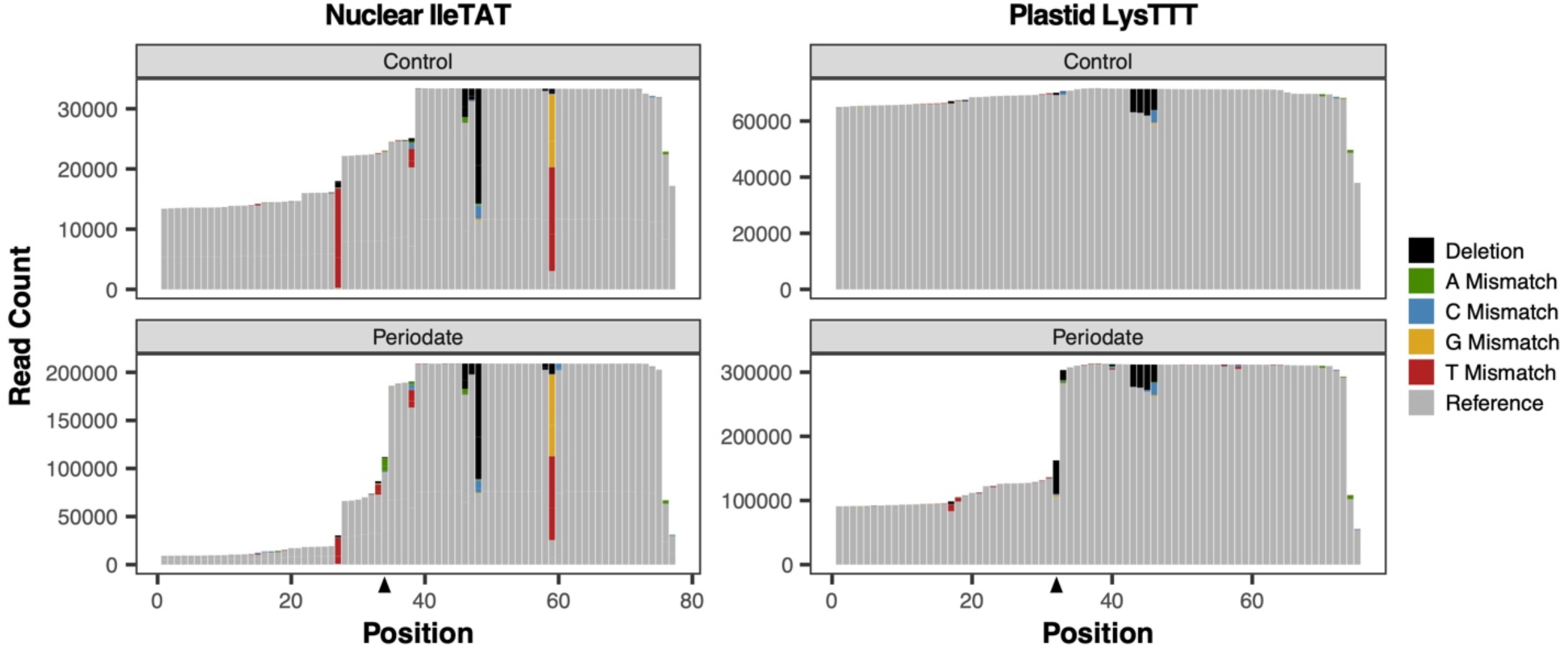
Read depth and misincorporation profiles for two tRNA isodecoder families that both show a major 5′ truncation point induced by periodate treatment. Position 34 in nuclear tRNA-IleTAT (left) and position 32 in plastid tRNA-LysTTT (right) – both of which correspond to position 33 in the Sprinzl coordinate system – show large drops in sequence coverage in periodate-treated samples (bottom) but not in control libraries (top). These positions are indicated by black triangles on the x-axes. In the reads that do cover these positions, there is also a large increase in the frequency of nucleotide misincorporations or deletions. Similar observations in other species (Katanski et al. 2022; Davidsen and Sullivan 2024) have been attributed to effects of periodate on 2-thio-modifications at the adjacent anticodon wobble position (34 in the Sprinzl coordinate system).

**Figure S8.**
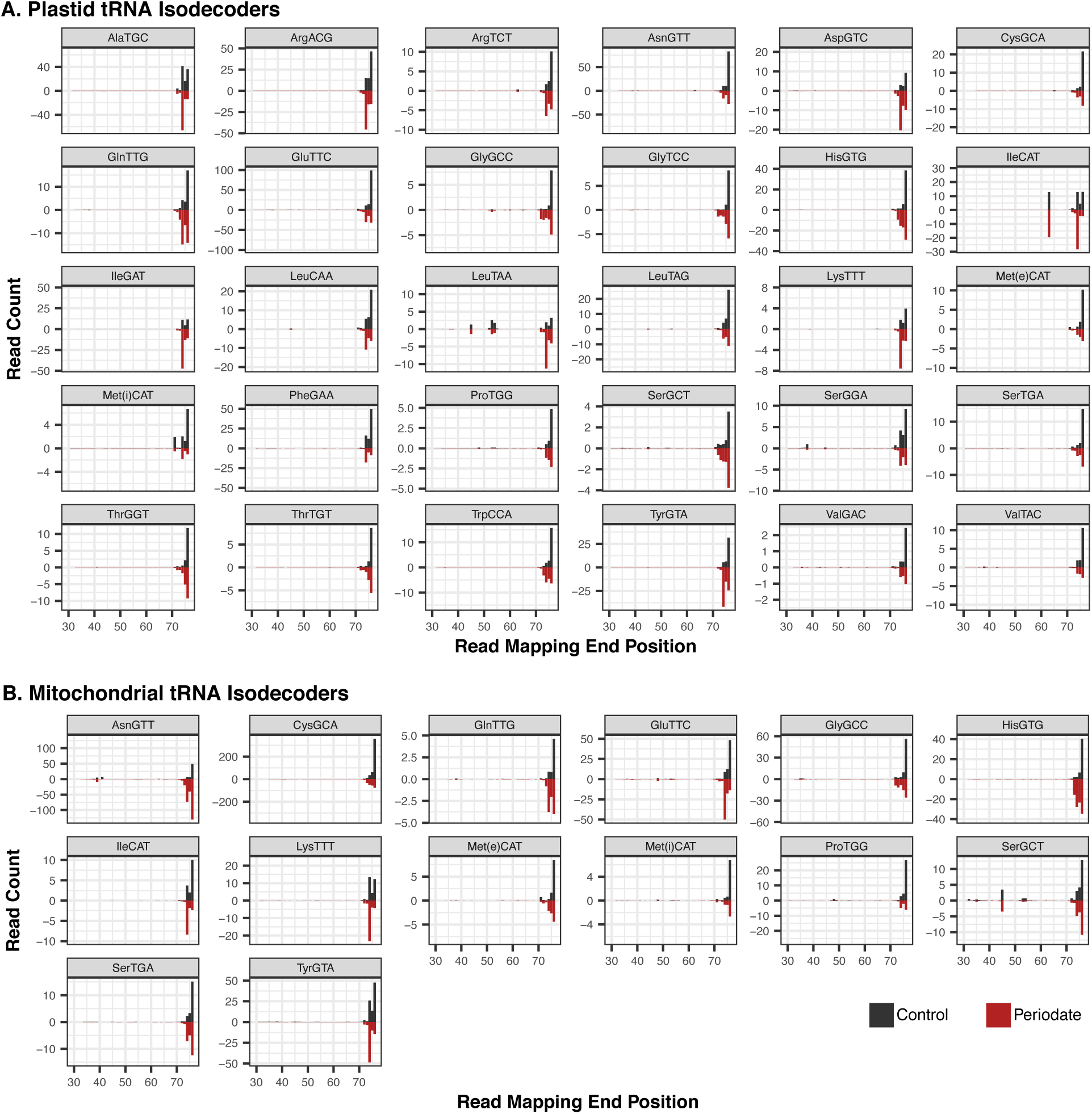
3′ mapping position of MSR-seq reads for *Arabidopsis* organellar tRNAs. Counts are represented per thousand reads that mapped to the (A) plastid or (B) mitochondrial tRNA gene set (averaged across the three biological replicates). Control libraries (black bars) are shown as positive values above the x-axis, while periodate-treated libraries (red bars) are shown as negative values below the y-axis. Mapping positions are standardized based on the Sprinzl coordinate system (Sprinzl et al. 1998). The mitochondrial AspGTC, SerGGA, and TrpCCA tRNAs are not included in this analysis because a large number of reads from their plastid-derived counterparts ambiguously map to these genes.

**Figure S9.**
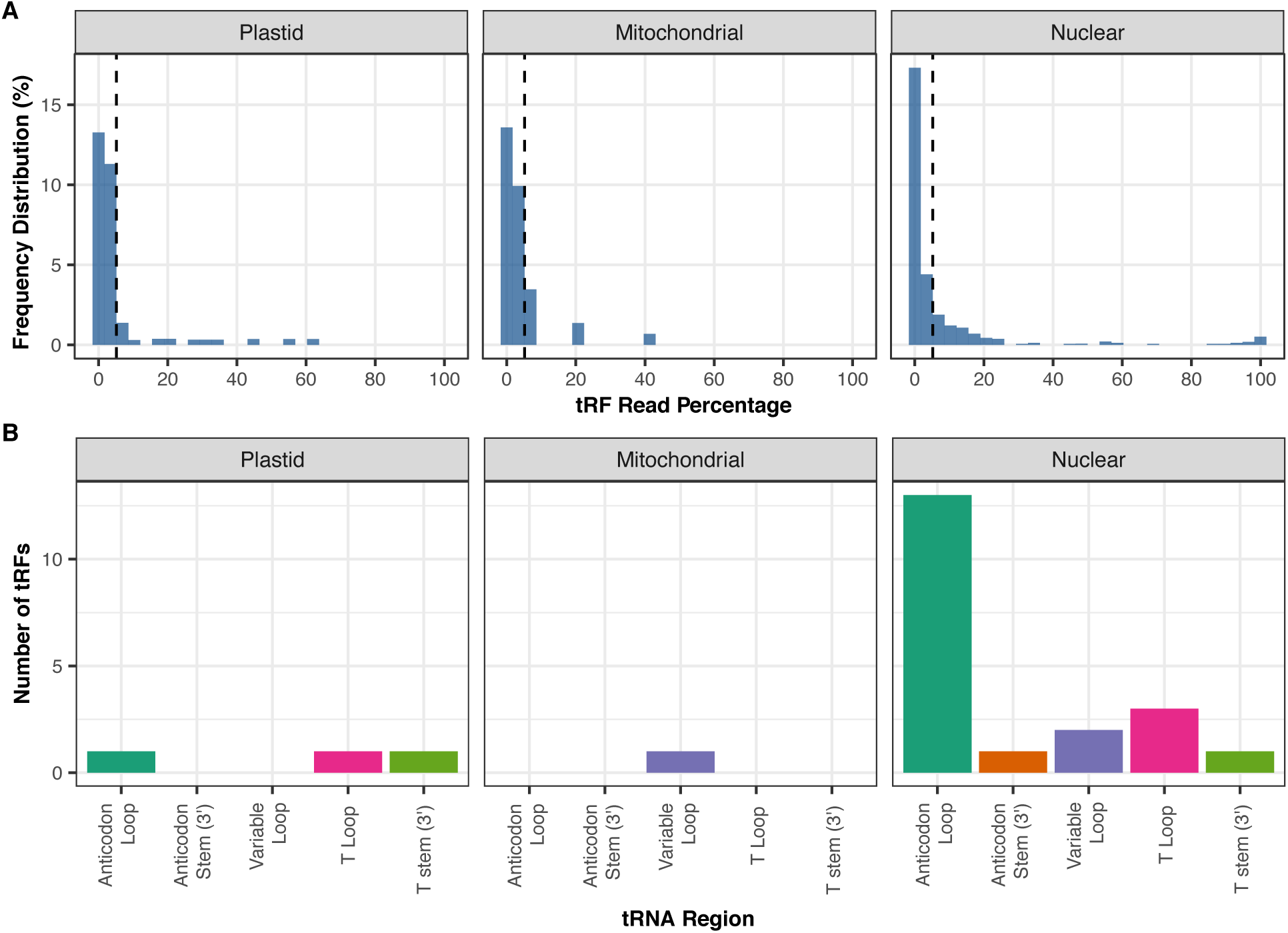
Summary of 5′ tRFs. (A) The proportion of sequence reads represented a minor fraction for most tRNA genes. A threshold of 5% (dashed line) was used to consider fragment a tRF for subsequent analysis. Analysis was limited to genes with a read count > 150. (B) Breakpoints were most often found in the Anticodon loop of nuclear tRNA genes.

**Figure S10.**
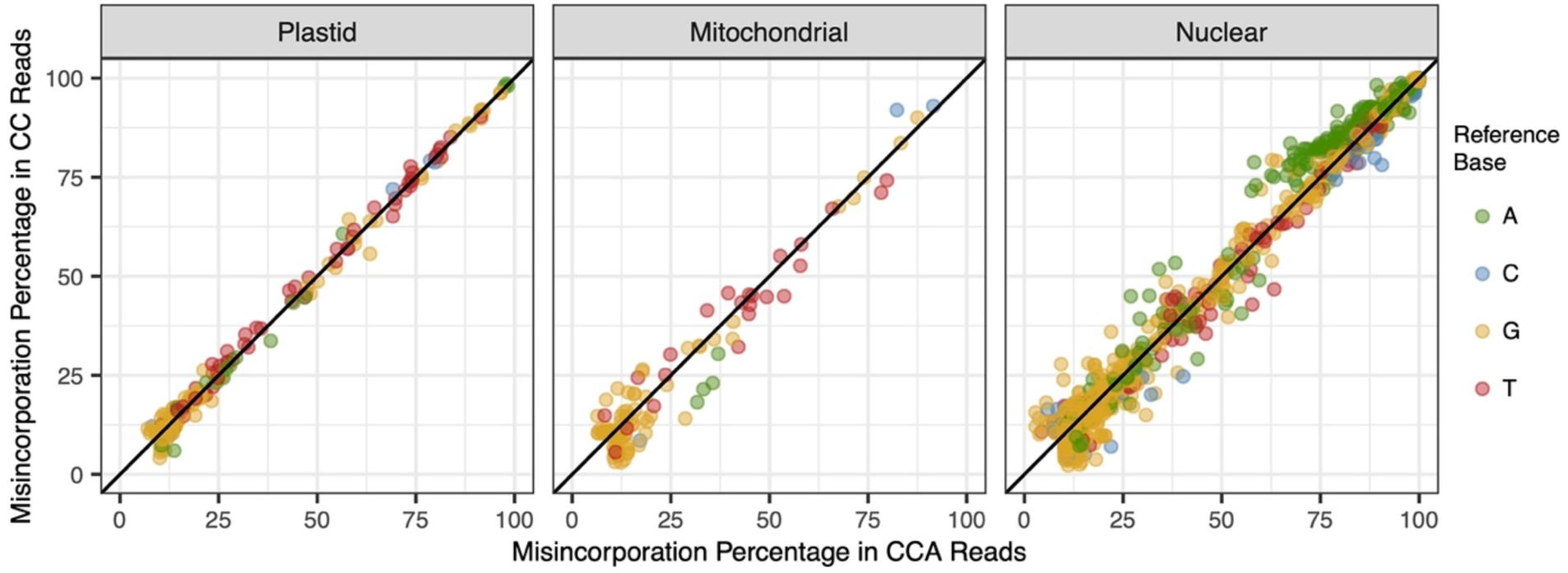
Relationship between nucleotide misincorporation rates in reads with intact CCA tails (aminoacylated tRNAs) vs. reads with CC tails (uncharged tRNAs). Each point represents the combined frequency of nucleotide substitutions and deletions at a specific position in one tRNA gene, averaged across three biological replicates that were treated with periodate. This value is calculated amongst the pool of reads with a full CCA tail (x-axis) or reads that lack their 3′ terminal nucleotide and end in CC (y-axis). Data are only reported for replicates with a read count >100 and for positions with a modification frequency of >10% in at least one sample. Reference tRNA genes are partitioned into panels by genome, and a one-to-one line is plotted in each panel. Misincorporation rates do not differ significantly between CCA and CC reads. As much, they do not provide evidence that aminoacylation levels are dependent on base modifications.

**Figure S11.**
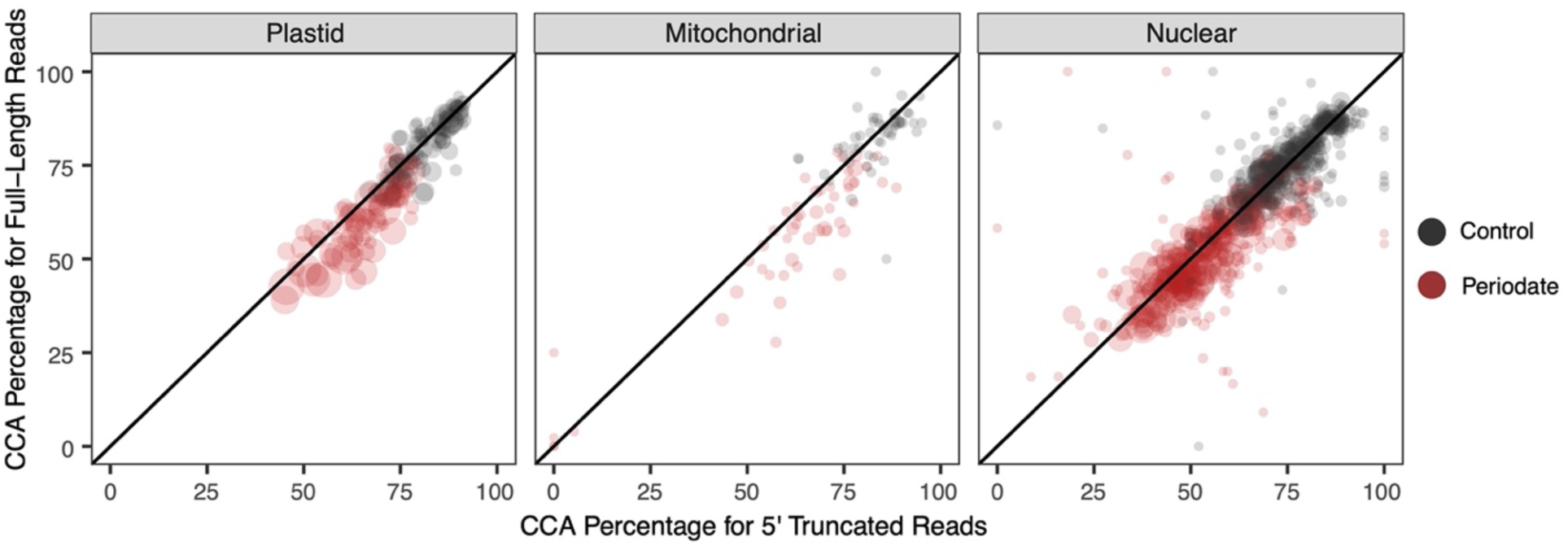
Relationship between percentage of reads with intact CCA tails and 5′ truncations as a proxy for “hard-stop” base modifications. Each points represents the percentage of reads with intact CCA tails after excluding reads that lacked more than just a single 3′ nucleotide. This value is calculated amongst the pool of reads that are truncated at their 5′ end by five or more nt (x-axis) or full-length reads (y-axis). Each point represents the combined frequency of nucleotide substitutions and deletions at a specific position in one tRNA gene, averaged across three biological replicates that were treated with periodate. Data are only reported for replicates with a (CC or CCA-containing) read count >150. Reference tRNA genes are partitioned into panels by genome, and a one-to-one line is plotted in each panel. In each of the three genome partitions, CCA percentages are significantly higher for 5′ truncated reads than for full-length reads in periodate-treated libraries (red points) but not in no-periodate controls (gray points). This observation is consistent with hypothesis that the base modifications that result in RT truncations affect tRNA function and increase aminoacylation rates/levels.

**Table S1.** Summary of MSR-seq libraries generated for this study, including number of reads produced, successfully processed (removal of expected adapter sequencing and merging R1 and R2 reads with BBMerge), and mapped the tRNA reference set.

| NCBI SRA | Experiment | RNA Extraction | Treatment | Rep | Read Pairs | Processed | Mapped |
| --- | --- | --- | --- | --- | --- | --- | --- |
| SRR30502882 | 1 | Acid-Phenol | No Periodate | 1 | 17664000 | 15080940 | 14110062 |
| SRR30502881 | 1 | Acid-Phenol | No Periodate | 2 | 14443230 | 12291168 | 11653350 |
| SRR30502880 | 1 | Acid-Phenol | No Periodate | 3 | 24655596 | 21714460 | 20546775 |
| SRR30502892 | 1 | Acid-Phenol | Periodate | 1 | 19364207 | 14956390 | 12953944 |
| SRR30502891 | 1 | Acid-Phenol | Periodate | 2 | 66433974 | 60449437 | 55430258 |
| SRR30502883 | 1 | Acid-Phenol | Periodate | 3 | 45130569 | 41270082 | 37775713 |
| SRR30502879 | 1 | Acid-Phenol | Predeacylated+Periodate | 1 | 49028388 | 44243871 | 39008946 |
| SRR30502878 | 1 | Acid-Phenol | Predeacylated+Periodate | 2 | 18369858 | 14175129 | 12367300 |
| SRR30502877 | 1 | Acid-Phenol | Predeacylated+Periodate | 3 | 33453924 | 28715989 | 25125094 |
| SRR30502887 | 2 | Acid-Phenol | No Periodate (with spike-in) | 1 | 4955018 | 4670581 | 4403102 |
| SRR30502886 | 2 | Acid-Phenol | No Periodate (with spike-in) | 2 | 2540908 | 2379755 | 2260925 |
| SRR30502876 | 2 | Acid-Phenol | Periodate (with spike-in) | 1 | 5552125 | 5136589 | 4642057 |
| SRR30502890 | 2 | Acid-Phenol | Periodate (with spike-in) | 2 | 5809426 | 5381792 | 4923051 |
| SRR30502885 | 2 | Trizol | No Periodate (with spike-in) | 1 | 3763956 | 3589193 | 3440449 |
| SRR30502884 | 2 | Trizol | No Periodate (with spike-in) | 2 | 4507211 | 4283316 | 3970242 |
| SRR30502889 | 2 | Trizol | Periodate (with spike-in) | 1 | 5848749 | 5507450 | 5065511 |
| SRR30502888 | 2 | Trizol | Periodate (with spike-in) | 2 | 4461382 | 4112669 | 3560700 |

**Table S2.**
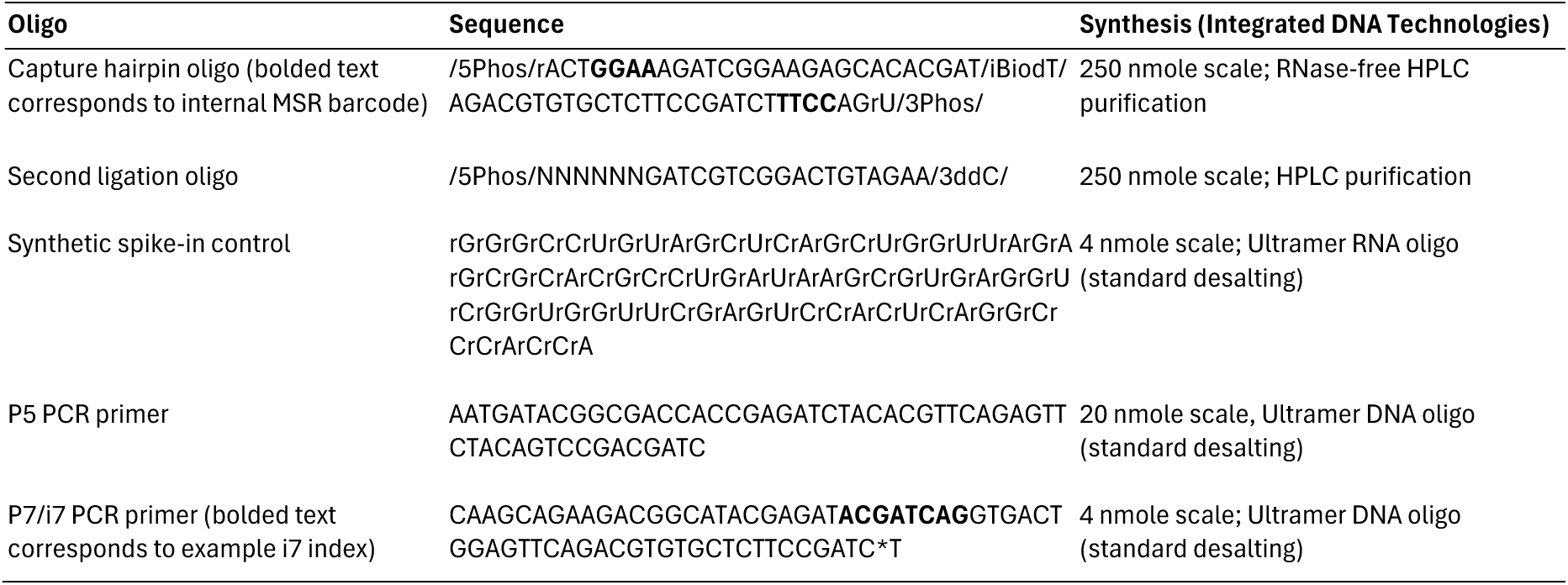
Oligonucleotides used in library construction

